# Constitutive TRIM22 expression within the respiratory tract identifies tissue-specific and cell-type dependent intrinsic immune barriers to influenza A virus infection

**DOI:** 10.1101/679159

**Authors:** Matthew Charman, Steven McFarlane, Joanna K. Wojtus, Elizabeth Sloan, Rebecca Dewar, Gail Leeming, Mohammed Al-Saadi, Laura Hunter, Miles Carroll, James P. Stewart, Paul Digard, Edward Hutchinson, Chris Boutell

**Affiliations:** MRC - University of Glasgow Centre for Virus Research, Glasgow, G61 1QH, UK; Division of Protective Immunity and Division of Cancer Pathobiology, Children’s Hospital of Philadelphia, Philadelphia, Pennsylvania, USA; The Roslin Institute, University of Edinburgh, Midlothian, EH25 9RG, UK; Institute of Infection and Global Health, University of Liverpool, Liverpool, L3 5RF, UK; National Infection Service, Public Health England, Porton Down, Salisbury, UK; University of Al-Qadisiyah, Al-Qadisiyah, Al-Diwaniyah, Iraq

## Abstract

We hypothesized that increased expression of antiviral host factors at portals of viral entry may protect exposed tissues from the constant threat of invading pathogens. Comparative transcriptomic analysis identified the broad-acting restriction factor TRIM22 (TRIpartite Motif 22) to be among the most abundantly expressed antiviral host factors in the lung, a major portal of entry for many respiratory pathogens. This was surprising, as TRIM22 is currently considered to be an interferon stimulated gene (ISG) product that confers protection following the activation of pathogen-induced cytokine-mediated innate immune defences. Using human respiratory cell lines and the airways of rhesus macaques, we experimentally confirmed high levels of constitutive TRIM22 expression in the lung. In contrast, TRIM22 expression in many widely used transformed cell lines could only be observed following immune stimulation. Endogenous levels of TRIM22 in non-transformed cells were sufficient to restrict human and avian influenza A virus (IAV) infection by inhibiting the onset of viral transcription independently of cytokine-mediated innate immune defences. Thus, TRIM22 confers a pre-existing (intrinsic) tissue-specific immune barrier to IAV infection in the respiratory tract. We investigated whether the constitutive expression of TRIM22 was a characteristic shared by other ISGs in human lung tissue. Transcriptomic analysis identified a large group of ISGs and IAV immuno-regulatory host factors that were similarly enriched in the lung relative to other mucosal tissues, but whose expression was downregulated in transformed cell-lines. We identify common networks of immune gene downregulation which correlated with enhanced permissivity of transformed cells to initiate IAV replication. Our data highlight the importance of tissue-specific and cell-type dependent patterns of pre-existing immune gene expression in the intrinsic intracellular restriction of IAV; findings highly relevant to the immune regulation of many clinically important respiratory pathogens.

**Author Summary:** The respiratory tract is a major portal of virus entry for many clinically important viruses, including seasonal and pandemic influenza A virus (IAV). We reasoned that cells within the respiratory tract might differentially express antiviral host factors to protect against the constant challenge of viral infection. We found the broad-acting antiviral protein TRIM22, conventionally regarded as an interferon stimulated gene (ISG) product upregulated in response to virus infection, to be constitutively expressed to high levels in the lung. We found that constitutive expression of TRIM22 restricted the initiation of human and avian IAV infection independently of cytokine-mediated innate immune defences. We identified pre-existing tissue-specific and cell-type dependent patterns of constitutive immune gene expression that strongly correlated with enhanced resistance to IAV replication from the outset of infection. Importantly, we show that these constitutive patterns of immune gene expression are lost or downregulated in many transformed cell lines widely used for respiratory virus research. Our data highlight the importance of pre-existing tissue-specific and cell-type dependent patterns of constitutive antiviral gene expression in the intracellular restriction of respiratory viral pathogens not captured in conventional cell culture model systems of infection.

## Introduction

Exposure to viral pathogens is a constant threat to all living things and vertebrates have evolved multiple lines of defence to suppress infection. If viruses succeed in penetrating non-specific barrier defences, the activation of pattern recognition receptors (PRRs) by pathogen- and damage-associated molecular patterns (PAMPs and DAMPs, respectively) leads to the activation of innate immune defences, culminating in the secretion of cytokines (including interferons) and the induction of hundreds of interferon stimulated gene (ISG) products [1–4]. ISGs include a wide range of antiviral effectors, and their induced expression from low basal levels to high functional levels plays an important role in limiting viral propagation to resolve pathogen infection [4–6]. However, the induction of this broad antiviral response necessitates pathogen detection by PRRs, which in the case of wild-type influenza A virus (IAV) requires the detection of aberrant viral RNAs (vRNAs) or defective interfering (DI) particles produced during virus replication for optimal induction [1, 3, 7–12]. Accordingly, delayed activation of innate immune defences provides a window of opportunity for viral pathogens to express immunosuppressive genes, which can inhibit or dampen the efficacy of host immune defences [13, 14]. A growing body of evidence suggests that this initial ‘gap’ in intracellular immunity is covered by intrinsic immunity, also known as intrinsic antiviral resistance or cell autonomous immunity [15–18].

Intrinsic immune effectors are constitutively expressed at levels sufficient to confer protection from the outset of infection. As a result, they can restrict the initiation or progress of viral replication prior to the pathogen-induced activation of PRRs and induction of innate immune defences [15, 16, 19–22]. Notably, many intrinsic antiviral host factors are themselves ISGs (intrinsically expressed ISGs; [18]), which can be further upregulated as a component of the innate immune response upon IFN production. Recent single-cell transcriptomic and reporter-assay studies have provided compelling evidence to support a biological role for intrinsic immunity during IAV infection both *in vitro* and *in vivo*. Cell culture studies have shown individual infected cells of the same lineage to be differentially permissive to IAV infection but to rarely induce the expression of IFN leading to the induction of ISGs [8, 11, 12, 23, 24]. Animal studies have shown lineage-specific patterns of IAV restriction that vary between cell-types, including lung epithelial, fibroblast, endothelial, and resident immune cells [25, 26]. Importantly, these lineage-specific patterns of restriction *in vivo* were shown to occur independently of IRF7 (interferon response factor 7), a critical transcriptional regulator of host innate immune defences to IAV infection [25, 27–32]. These data suggest that intrinsic patterns of constitutive host gene expression are likely to play an important role in limiting IAV replication immediately upon pathogen entry into susceptible host cells, thereby reducing the need to prematurely activate potentially harmful pro-inflammatory innate immune defences [2, 33, 34]. However, evidence for tissue-specific or cell-type intrinsic immune effectors that may restrict IAV replication has remained lacking.

We hypothesized that localized intrinsic immune barriers might exist due to tissue-specific patterns of gene expression at common portals of viral entry. We tested this hypothesis in cells derived from the respiratory tract, a major portal of virus entry for many clinically important pathogens, including seasonal and pandemic influenza viruses [35]. In order to identify antiviral genes that might be differentially expressed in the respiratory tract, we initially focused on TRIpartite Motif (TRIM) proteins, a family of over 70 members that participate in a wide range of cellular processes, including multiple aspects of immune regulation and antiviral defence [36–39]. Many TRIM proteins are strongly upregulated in response to IFN signalling and are well established to act as regulators or effectors of innate immunity during virus infection [5, 36–38, 40–45]. Other TRIM proteins are constitutively expressed and known to directly mediate intrinsic immune defences [16, 17, 19, 46–49], including TRIM32 and TRIM41 which have been reported to restrict IAV replication through the targeted degradation of PB1 (polymerase basic protein 1) and NP (nucleoprotein), respectively [21, 22].

We focussed particularly on TRIM22, which we identified to be amongst the most abundantly expressed TRIM family members in the respiratory tract. TRIM22 has been implicated in the cellular restriction of a broad range of viruses including encephalomyocarditis virus (EMCV), hepatitis B virus (HBV), hepatitis C virus (HCV), human immunodeficiency virus (HIV), and IAV [50–55]. Studies in transformed cultured cells and primary lymphocytes have shown TRIM22 to be an ISG, strongly upregulated by immune stimuli including type-I (α, β) and -II (γ) IFNs; interleukins (IL-1β, -2 and -15); progesterone and tumour necrosis factor-α (TNF-α) [54–56]. Accordingly, TRIM22 has been shown to inhibit viral infection following its induced expression as an ISG by restricting the onset of viral transcription or by targeted degradation of viral proteins [50-54, 57-60]. With respect to IAV, transient transfection studies in transformed cells have shown TRIM22 to mediate the ubiquitination and proteasome-dependent degradation of NP [55, 61], and to restrict IAV propagation as an effector of the type-I IFN response [55].

Here, we show that TRIM22 is constitutively expressed to high levels in the respiratory tract and non-transformed cells of lung origin independently of immune stimulus or viral infection. We demonstrate that the endogenous levels of TRIM22 expression are sufficient to restrict human and avian IAV infection by inhibiting the onset of viral transcription independently of cytokine-mediated innate immune defences. Thus, we identify TRIM22 to confer a pre-existing (intrinsic) intracellular immune barrier to IAV infection within cells of the respiratory airway. Consistent with our hypothesis, these high levels of TRIM22 expression are not present in all cell-types or tissues. Equally importantly, transcriptomic analysis revealed TRIM22 to be amongst a large group of IAV immune regulators that are downregulated in transformed cells which share common networks of immune system disruption correlating with enhanced permissivity to IAV replication. Collectively, our data demonstrate that tissue-specific and cell-type dependent patterns of pre-existing immune gene expression to play a critical role in the intrinsic intracellular restriction of IAV from the outset of infection. These findings are highly relevant to the immune regulation of many clinically important respiratory pathogens.

## Results

### TRIM22 is constitutively expressed at high levels in the respiratory tract independently of immune stimulus or virus infection

As the respiratory tract is a major portal for virus entry, we hypothesized that cells in the respiratory mucosa might express antiviral proteins to higher levels than cells in less exposed locations, thereby creating localized pre-existing (intrinsic) immune barriers to virus infection. We initially explored this hypothesis using RNA-seq data and protein expression records from Human Protein Atlas (HPA; https://www.proteinatlas.org; [62, 63]) and Genotype-Tissue Expression (GTEx) project (https://gtexportal.org/home/; [64]). We focussed on the TRIM family of proteins, as many members of this family are known to directly or indirectly mediate antiviral immune responses to a wide range of viruses [37]. Of the TRIM family, RNA-seq data from HPA indicated that three members (TRIM8, 22, and 28) had the highest transcript expression levels in human lung tissue (Fig 1A, red circles; S1A Table), with TRIM22 being the most abundantly expressed. Notably, expression of these TRIMs was substantially higher than that of TRIM32 and TRIM41, two previously identified intrinsic antiviral regulators of IAV (Fig 1A, blue circles; [21, 22]). TRIM22 transcript levels were most abundantly expressed in the spleen, lymph node, appendix, gallbladder, and lung (Fig 1B, TRIM22 coloured circles; S1B Table), with expression values exceeding the 95% confidence interval for median TRIM22 expression across all tissues. In contrast, transcript levels for both TRIM32 and TRIM41 in the lung were close to their respective median tissue expression values (Fig 1B, TRIM32/41 red circles; S1B Table). Analysis of RNA-seq data obtained by the GTEx project independently confirmed TRIM8, 22, and 28 to be the most abundantly expressed TRIMs in human lung tissue (S1 Fig, S1A Table), with the highest levels of TRIM22 expression observed in the spleen and lung (S1 Fig, S1C Table). Together, these data demonstrate that TRIM22 transcript levels show tissue-specific patterns of gene expression and to be enriched within the lung relative to other tissues or TRIM family members. Analysis of HPA immunohistochemistry (IHC) expression records demonstrated the nasopharynx and bronchus to be among tissues with the highest levels of TRIM22 expression (Fig 1C, D). These relatively high levels of tissue-specific protein expression suggest that TRIM22 could make a substantial contribution to a pre-existing and localized intrinsic immune barrier to respiratory airway infection.

**FIG 1.**
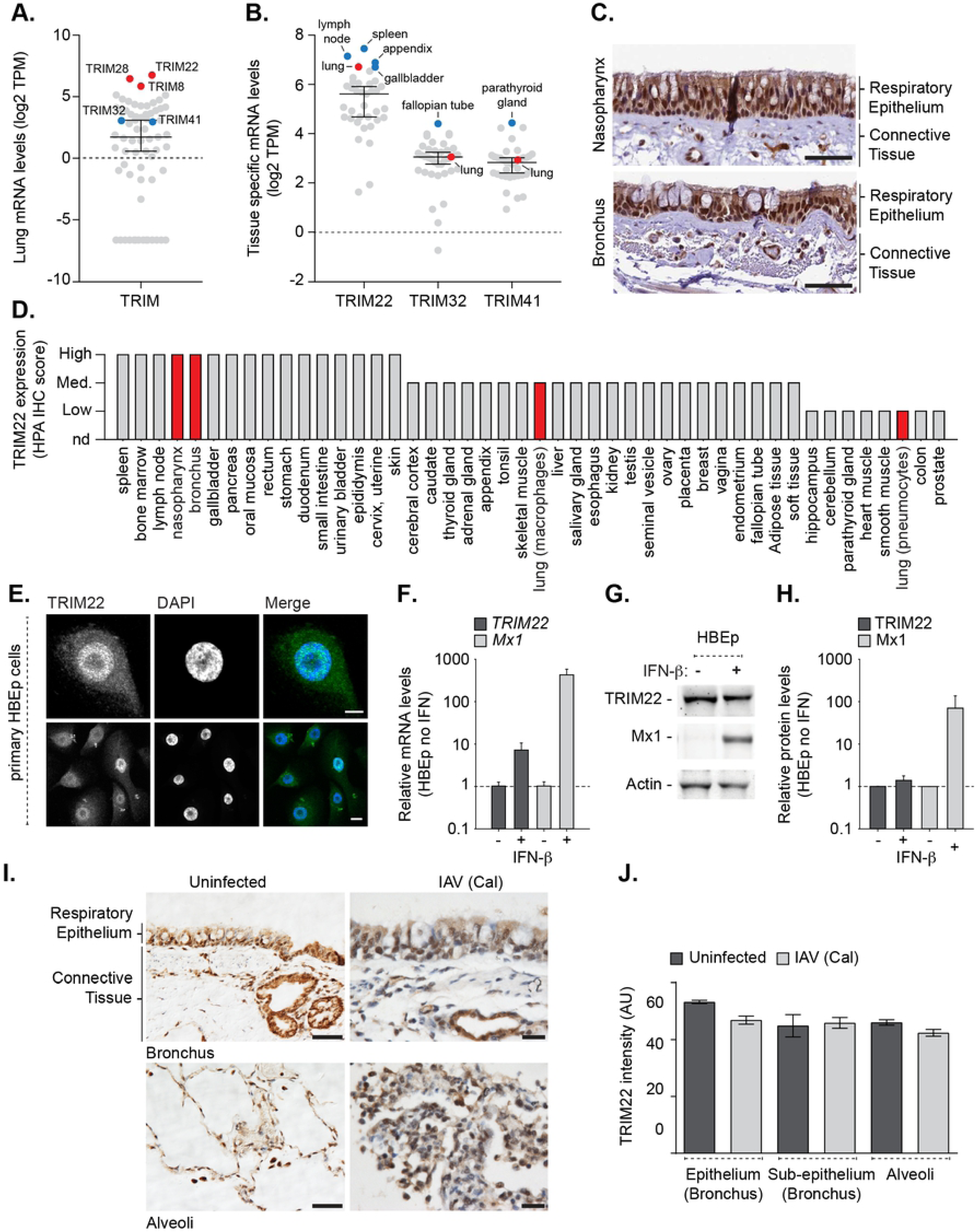
TRIM22 is constitutively expressed in the respiratory tract independently of IAV infection. (A) mRNA transcript levels (log2 transcripts per million; TPM) of TRIM family members in human lung tissue (S1A Table). Black line: median TRIM transcript expression; whisker: 5^th^ to 95^th^ percentile range. (B) TRIM22, TRIM32, and TRIM41 transcript levels across a range of human tissues (S1B Table). Black line: median; whisker: 5^th^ to 95^th^ percentile range. (C) Histological sections of human respiratory epithelium from the upper (nasopharynx; patient ID 3624) and lower (bronchus; patient ID 3987) airway. TRIM22 is labelled by immunohistochemistry (IHC; brown) and tissue counterstained with haematoxylin and eosin. Scale bars are 50 µm. (D) Quantitation of TRIM IHC staining across a range of tissue (as indicated). Red bars: Quantitation of TRIM22 in lung tissue sections. (A-D) Data adapted under creative commons license from the Human Protein Atlas (HPA; https://www.proteinatlas.org; [62, 63]). (E) Confocal micrographs of primary human bronchial epithelial (HBEp) cells stained for TRIM22 by indirect immunofluorescence (green). Nuclei stained with DAPI (blue). Scale bars; 10 µm (top panel) and 20 µm (bottom panel). (F-H) Primary HBEp cells were treated with (+) or without (-) IFN-β (100 IU/ml) for 24 h. (F) Western blots of HBEp whole cell lysates (WCLs) probed for TRIM22 and Mx1 expression. Actin is shown as a loading control. (G) Quantitation of western blots (as shown in F). Values normalized to actin and expressed relative to no IFN treatment; n=3, means and standard deviation (SD) shown. (H) qRT-PCR for *TRIM22* and *Mx1* mRNA transcript levels in control or IFN treated HBEp cells. Values normalized to no IFN treatment; n=3, means and SD shown. (I) Histological sections of uninfected or influenza A virus (IAV; A/California/04/09(H1N1), Cal) infected cynomolgus macaque respiratory epithelium from the bronchus and alveoli (as indicated). TRIM22 is labelled by IHC (brown) and tissue counterstained with haematoxylin. Scale bars; 50 and 20 µm (left and right panels, respectively). (J) Automated quantitation of TRIM22 IHC staining in uninfected or infected cynomolgus macaque respiratory tissue from whole-slide scans. Means and SD from four uninfected and three infected animals are shown.

However, the above data contrast with many previous studies of transformed cultured cells and primary lymphocytes, in which TRIM22 expression is strongly upregulated upon viral infection or immune stimulation [54–56]. To resolve this discrepancy, we first examined how TRIM22 expression in primary human bronchial epithelial (HBEp) cells responded to IFN stimulation. Using a validated TRIM22 antibody (S2 Fig), TRIM22 was readily detectable in unstimulated cells by immunofluorescence (Fig 1E), showing the same predominantly nuclear localisation observed by IHC in respiratory epithelia (Fig 1C). The addition of IFN-β caused an intense upregulation of the ISG Mx1 at both the transcript and protein level. In contrast, TRIM22 expression in the same cells was only increased at the transcript level and not detectably increased at the protein level (Fig 1F-H). We conclude that TRIM22 is constitutively expressed to high levels in non-transformed respiratory epithelial cells independently of immune stimulus. Next, we analysed how TRIM22 expression in the respiratory tract responded to viral infection. As mice lack an orthologue to human TRIM22 (https://www.ncbi.nlm.nih.gov/homologene/?term=trim22), we examined tissue from cynomolgus macaques (*Macaca fascicularis*), whose TRIM22 has 92% amino acid identity to human TRIM22. As in human tissue (Fig 1C, D), TRIM22 was constitutively expressed in the respiratory tract of uninfected macaques, specifically in epithelia of the airways, sub-mucosal glands, alveoli, and in alveolar macrophages, with little staining observed in the sub-epithelial connective tissue (Fig 1I, uninfected). In order to determine whether TRIM22 expression increased during infection, we compared healthy macaques with those infected with IAV (A/California/04/2009 (H1N1); Cal). These macaques had previously been shown to be infected and to be undergoing an induced innate immune response at the point of euthanasia [65]. Automated staining and quantitation of sectioned samples demonstrated TRIM22 expression did not increase in the respiratory tract of IAV infected macaques (Fig 1I, J). In contrast to its expression as an ISG in other settings [54, 55], these data demonstrate that the high levels of constitutive TRIM22 expression observed in primary HBEp cells is representative of its expression profile in the epithelium of the respiratory tract independently of immune stimulation.

### Constitutive TRIM22 expression correlates with low permissivity to IAV infection

We next wished to investigate the antiviral properties of constitutively expressed TRIM22. However, while we could demonstrate strong constitutive expression of TRIM22 in HBEp cells, these primary cells are challenging to maintain at high densities for functional studies due to the rapid onset of cellular senescence. Accordingly, to identify a tractable cell line that maintained constitutive expression of TRIM22, we screened a panel of human cell lines for TRIM22 transcript and protein expression levels with and without IFN-β stimulation. In all virally-transformed cell lines examined (HEK 293T, HeLa, HEp2, and A549), TRIM22 was either absent or, as observed for the ISG Mx1, only detected following IFN-β stimulation (Fig 2A-D, S3 Fig). Thus, IAV infection studies that have utilize transformed cell lines would not capture the endogenous antiviral properties of TRIM22 observed at the site of natural infection (Fig 1; [55, 61]). In contrast, both primary and human telomerase reverse transcriptase (hTERT)-immortalized human lung fibroblasts (MRC5 and MRC5t, respectively) retained constitutive expression of TRIM22 independently of IFN-β stimulation (Fig 2A-D, S3 Fig). As single-cell transcriptomics experiments have shown lung fibroblasts to be susceptible to IAV infection *in vivo* [25], we chose MRC5t cells as a model cell line to study the effects of constitutive TRIM22 expression on respiratory virus infection.

**FIG 2.**
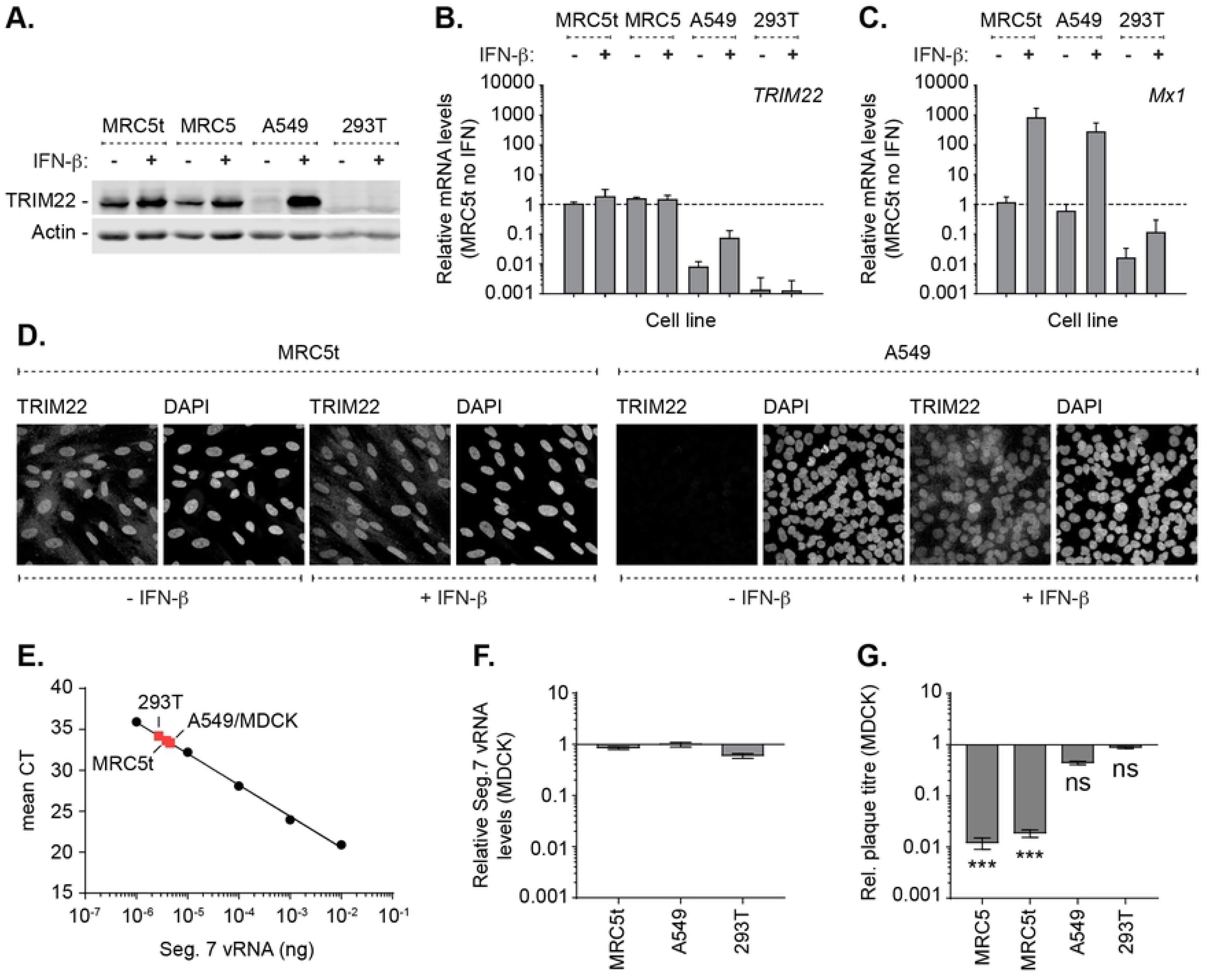
TRIM22 is constitutively expressed in human diploid lung cells. (A-C) Primary and hTERT immortalized human lung fibroblast (MRC5 and MRC5t, respectively), human lung adenocarcinoma epithelial (A549), and SV40-transformed human kidney epithelial (HEK 293T; 293T) cells were treated with (+) or without (-) IFN-β (100 IU/ml) for 24 h. (A) Western blots of WCLs probed for TRIM22 expression. Actin is shown as a loading control. (B/C) qRT-PCR quantitation of *TRIM22* and *Mx1* mRNA transcript expression levels, respectively. Values normalized to MRC5t cells without IFN treatment; n=3, means and SD shown. (D) Confocal micrographs of MRC5t and A549 cells with or without IFN treatment (as described in A). TRIM22 labelled by indirect immunofluorescence. Nuclei stained with DAPI. (E) MDCK, 293T, A549, and MRC5t cells were infection with IAV (A/Puerto Rico/8/1934(H1N1), PR8) at a MOI of 1 PFU/cell (based on MDCK cell titres) in the presence of cycloheximide at 2 h prior to nuclear extraction and RNA isolation. qRT-PCR quantitation of PR8 segment 7 (seg. 7) viral RNA (vRNA) levels in isolated nuclei. Black circles: synthetic seg. 7 vRNA loading control standards (ng); Black line: semilog non-linear regression (R^2^ = 0.99); Red squares: seg. 7 vRNA levels detected in the nuclei of infected cells (as indicated). (F) Seg. 7 vRNA levels (as shown in E) expressed relative to vRNA levels isolated from infected MDCK nuclei. (E/F) n=3, means and SD shown. (G) Cell monolayers were infected with serial dilutions of IAV (A/WSN/1933(H1N1), WSN). Plaque numbers in each cell line were expressed relative (rel.) to MDCK cells (rel. plaque titre); n=3, means and SD shown. One-way ANOVA Kruskal-Wallis test (*** *P* < 0.001; ns, not significant).

In order to investigate whether patterns of TRIM22 expression correlated with permissivity to IAV infection, we carried out IAV infections in a panel of cells with different TRIM22 expression phenotypes: constitutive (MRC5 and MRC5t), interferon-inducible (A549), or absent (HEK 293T). First, to determine if constitutive TRIM22 expression correlated with a block to viral entry, cells were infected with IAV (A/Puerto Rico/8/1934 (H1N1); PR8) at a multiplicity of infection (MOI) of 1 PFU/cell (based on titres derived in MDCK cells) in the presence of cycloheximide to prevent viral protein synthesis and genome replication. Similar levels of genome segment 7 (Seg. 7) vRNA were detected in the nuclei of infected cells at 2 hours post-infection (hpi) independently of cell lineage, indicating that all cells were equally permissive to viral entry and genome translocation to the nucleus (Fig 2E, F). Next, we tested whether TRIM22 expression patterns correlated with an inhibition of viral replication. To do this we compared the plaque titre of IAV (A/WSN/33 (H1N1); WSN) in each cell type to that in MDCK cells. IAV formed plaques in A549 and HEK 293T cells with an efficiency approximately equal to that in MDCK cells. In contrast, plaque formation in MRC5 and MRC5t cells was strongly suppressed (80 to 100-fold relative to MDCK cells; Fig 2G). The correlation of constitutive TRIM22 expression with restricted IAV plaque formation in both MRC5 and MRC5t cells suggested that constitutive TRIM22 expression may confer a pre-existing (intrinsic) intracellular immune barrier to IAV replication.

### Constitutively expressed TRIM22 restricts the initiation of IAV replication

In order to examine in more detail how TRIM22 expression patterns influenced permissivity to IAV infection, we focussed on MRC5t cells, in which TRIM22 is constitutively expressed, and A549 cells, in which TRIM22 is an ISG (Fig 2A). First, we compared patterns of TRIM22 expression during IAV (WSN) infection. In MRC5t cells, TRIM22 showed similar levels of constitutive expression both before and during a single-cycle of infection (up to 8 hpi; MOI of 1 PFU/cell based on MRC5t titres). In contrast, TRIM22 was barely detectable in A549 cells under equivalent infection conditions, as was the ISG Mx1 (Fig 3A). TRIM22 and Mx1 were only detectable in A549 cells following multi-cycle replication (48 hpi; MOI of 0.01 PFU/cell based on A549 titres), indicative of the induction of innate immune defences and ISG expression in response to PAMPs (aberrant vRNAs and DI particles) produced during viral replication (Fig 3B; [1, 11]).

**FIG 3.**
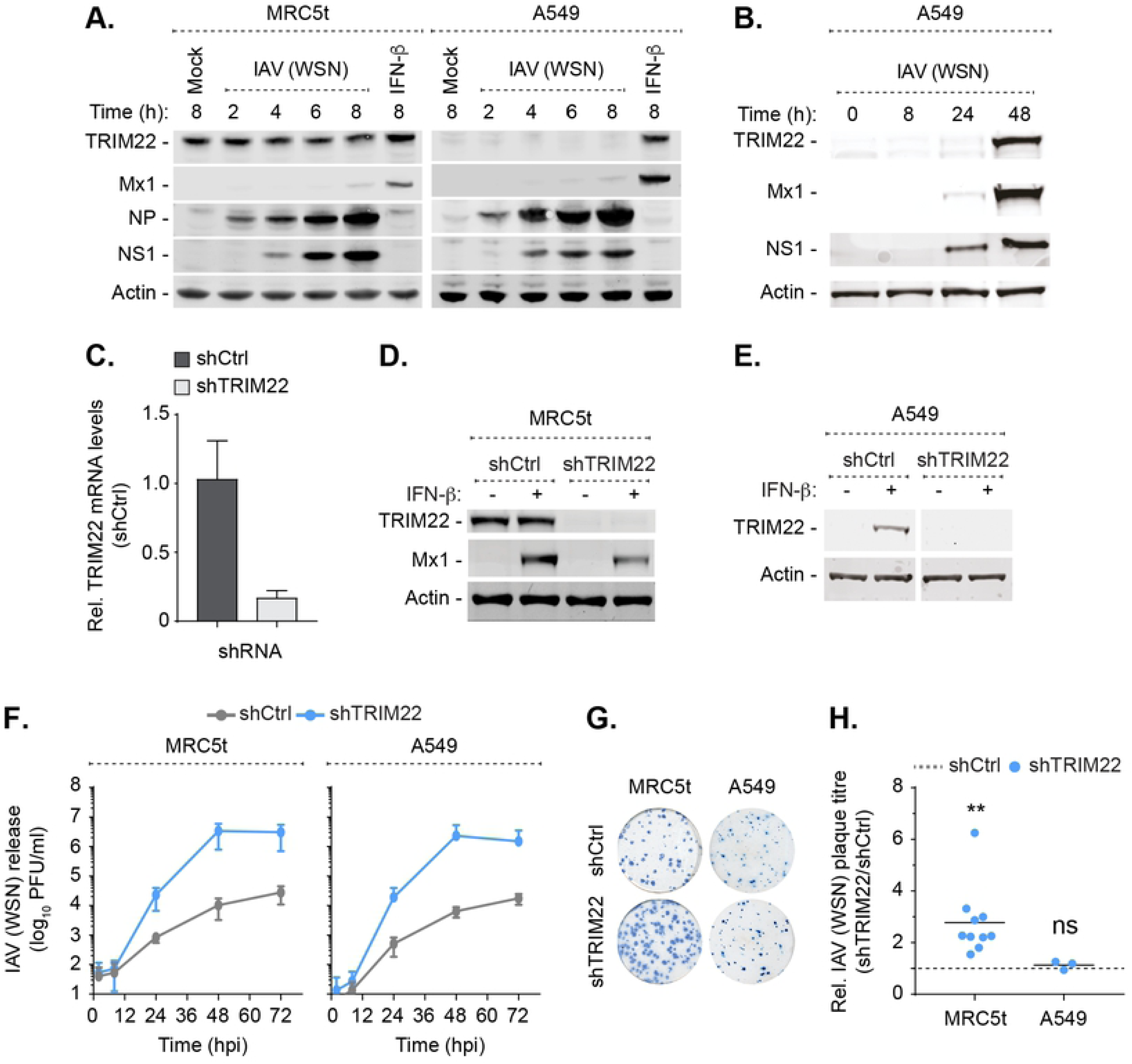
Constitutive TRIM22 expression confers intrinsic antiviral immunity. (A) MRC5t and A549 cells were mock-treated, IFN-β (100 IU/ml) stimulated or infected with IAV (A/WSN/1933(H1N1), WSN) at a MOI of 1 PFU/cell (based on MDCK titres) for the indicated times (h). Western blots of infected or treated WCLs (as indicated) probed for TRIM22, Mx1, and viral protein (NP and NS1) expression. Actin is shown as a loading control. (B) A549 cells were infected with IAV (WSN; MOI of 0.01 PFU/cell based on MDCK titres) and harvested at the indicated times prior to western blotting (as in A). (C-E) MRC5t and A549 cells were stably transduced to express non-targeting control (shCtrl) or TRIM22-targeting (shTRIM22) shRNAs. (C) qRT-PCR quantitation of *TRIM22* mRNA levels in MRC5t shCtrl and shTRIM22 cells. Values normalized to shCtrl; n=3, means and SD shown. (D) MRC5t shCtrl and shTRIM22 cells were treated with (+) or without (-) IFN-β (100 IU/ml) for 24 h. WCLs were analysed by western blotting for TRIM22 and Mx1. Actin is shown as a loading control. (E) Western blot analysis of A549 shCtrl and shTRIM22 treated cells (as in D). (F) MRC5t and A549 shCtrl and shTRIM22 cells were infected with IAV (WSN) at 0.001 PFU/cell (based on parental cell line titres). Media were harvested at the indicated time points and IAV plaque titres determined on MDCK cells; n=3, means and SD shown. (G) Representative immunocytochemistry images of IAV plaque formation (NP staining) in MRC5t and A549 infected shCtrl and shTRIM22 cell monolayers (50-100 PFU/monolayer based on parental cell line titres). (H) Relative (rel.) IAV plaque titre (plaque titre in shTRIM22 cells / plaque titre in shCtrl cells) in MRC5t and A549 infected cell monolayers. All data points shown; line: mean. One-sample two-tailed t test (hypothetical mean of 1; ** *P* < 0.005; ns, not significant).

Next, in order to identify the specific effects of TRIM22 we generated stable MRC5t and A549 cell lines expressing non-targeting control and TRIM22-targeting short hairpin RNAs (shCtrl and shTRIM22, respectively). The expression of *TRIM22* mRNA in MRC5t shTRIM22 cells was substantially depleted relative to control cells (Fig 3C, S2 Fig). As a result, the expression of TRIM22 protein was significantly knocked down in both MRC5t and A549 cells, with or without IFN-β stimulation (Fig 3D, E). To identify the contribution of TRIM22 to the cellular restriction of IAV replication, we infected these cells with IAV (WSN) at a low MOI (0.001 PFU/cell based on titres relative to each parental cell line) and used MDCK plaque assays to measure the release of infectious virus into the growth media over time. Depletion of TRIM22 enhanced IAV replication in both MRC5t and A549 cells relative to their respective controls, confirming the ability of TRIM22 to restrict IAV replication (Fig 3F; [55]); a phenotype attributable in A549 cells to the ISG induction of TRIM22 during multi-cycle replication (Fig 3B; [55]). We next examined the relative plaque titre of IAV in TRIM22 depleted cells compared to that in their respective control cell lines, reasoning that the ability of the virus to form plaques in these cells would reflect the ability of constitutively expressed TRIM22 to restrict IAV replication from the outset of infection. We infected cells with serial dilutions of IAV (WSN) and counted the number of plaques formed by 36 hpi (Fig 3G). In MRC5t cells, depletion of TRIM22 significantly enhanced the plaque titre of IAV. In contrast, depletion of TRIM22 in A549 cells had no significant effect (Fig 3H). We conclude that the constitutive expression of TRIM22 in non-transformed respiratory cells confers a pre-existing immune barrier to IAV infection that restricts viral replication leading to plaque formation.

### Constitutively expressed TRIM22 provides broad protection against IAV independently of the IFN response

Even though TRIM22 is constitutively expressed in MRC5t cells, it could act to restrict IAV infection by modulating innate immune signalling pathways, a widespread mode of action among the TRIM family [37, 45]. If this was the case, TRIM22 might be required to potentiate an innate antiviral immune response, but not provide a direct intracellular barrier to infection itself. To examine this, we tested the effects of TRIM22 in the presence of the Janus associated kinase (JAK) inhibitor Ruxolitinib (Ruxo), which has been shown to inhibit the induction of cytokine-mediated innate immune defences and ISG expression in response to IAV infection [66]. We first determined a concentration of Ruxo that would block ISG induction (*Mx1* and *ISG15*) in MRC5t cells following IFN-β treatment (Fig 4A; 4 µM). Comparing the effects of Ruxo (4 μM) or DMSO (carrier control) treatment on the relative plaque titre of IAV in TRIM22 depleted or control MRC5t cells showed that TRIM22 depletion caused the same increase in relative plaque titre regardless of the inhibition of JAK-STAT signalling (Fig 4B). Thus, constitutively expressed TRIM22 restricts IAV replication independently of pathogen-induced cytokine-mediated innate immune defences.

**FIG 4.**
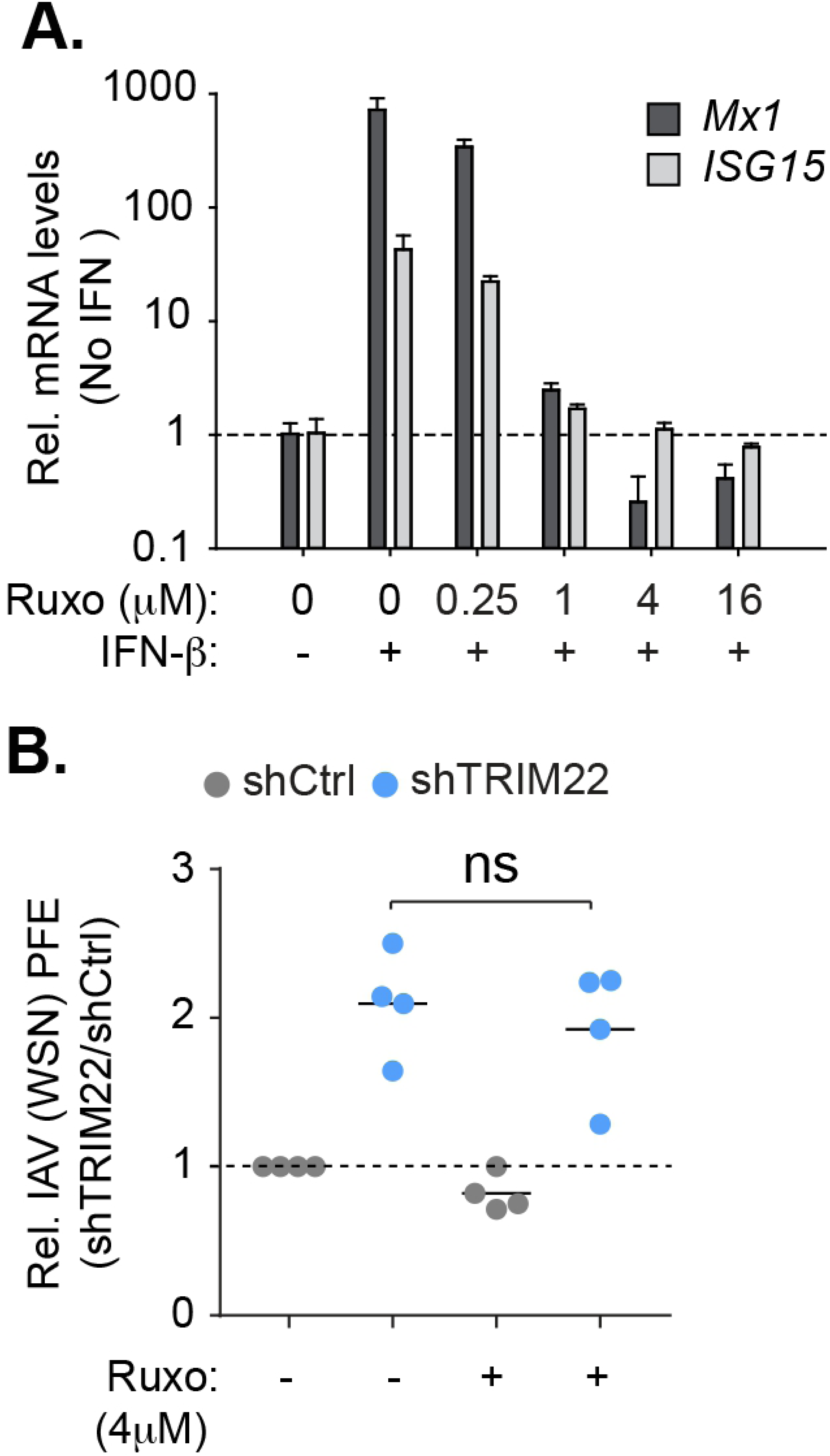
Constitutive TRIM22 expression restricts IAV infection independently of the induction of cytokine-mediated innate immune defences. (A) MRC5t cells were pre-treated for 1 h with increasing concentrations (µM) of Ruxolitinib (Ruxo) or DMSO (carrier control) prior to stimulation with (+) or without (-) IFN-β (100 IU/ml) for 24 h (in the presence or absence of drug, as indicated). qRT-PCR quantitation of ISG (*Mx1* and *ISG15*) mRNA levels from RNA extracted from treated cells. Values normalized to DMSO-only treatment; RQ and RQmin/max plotted. (B) Quantitation of relative (rel.) IAV (WSN) plaque titre (# plaques shTRIM22/# plaques shCtrl) in MRC5t cell monolayers treated with Ruxo (4 µM) or DMSO (as in A). Values normalized to infected shCtrl cell monolayers treated with DMSO per experiment. All data points shown; line: mean. One-sample two-tailed t test (hypothetical mean of 1; ** *P* < 0.005; ns, not significant).

As WSN is a highly laboratory-adapted strain, we tested whether constitutively expressed TRIM22 was effective against other IAV strains using an immunofluorescent focus-forming assay to measure the proportion of IAV NP positive cells at 8 hpi. Depletion of TRIM22 increased the focus-forming efficiency of a panel of influenza A viruses, including two avian strains (Fig 5A, B). Thus, constitutively expressed TRIM22 provides a broad-acting intrinsic immune barrier to IAV that restricts NP expression, protecting non-transformed respiratory cells against human and avian IAV from the outset of infection.

**FIG 5.**
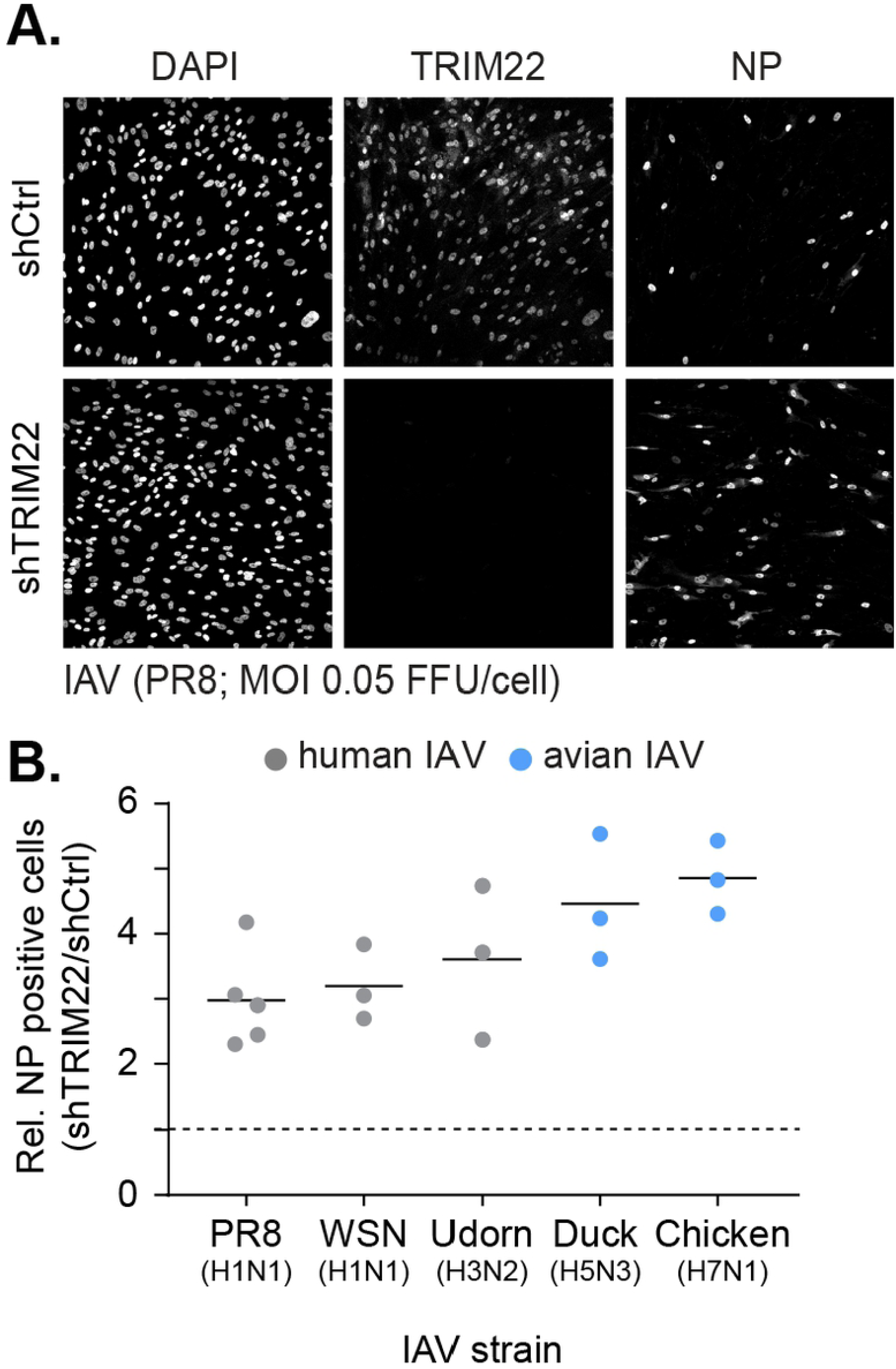
Constitutive TRIM22 expression restricts the initiation of IAV infection in a strain independent manner. MRC5t shCtrl or shTRIM22 cells were individually infected with a panel of human (WSN, PR8, and A/Udorn/307/1972(H3N2); Udorn) or avian (A/Duck/Singapore/5/1997(H5N3); Duck and A/Chicken/Italy/1067/1999(H7N1); Chicken) IAVs at a MOI of 0.05 FFU/cell (based on MRC5t titres) for 8h. (A) Representative confocal micrographs of PR8 infected MRC5t shCtrl or shTRIM22 cells. TRIM22 and IAV NP labelled by indirect immunofluorescence. Nuclei stained with DAPI. (B) Relative (Rel.) fold increase in NP antigen positive cells (shTRIM22/shCtrl) in IAV infected MRC5t cell monolayers. n ≥ 3, all data points shown; line: mean.

### TRIM22 provides intrinsic immunity against IAV by limiting the onset of viral transcription

Having established that the constitutive expression of TRIM22 confers a pre-existing intrinsic immune barrier to IAV, we wished to determine the point in viral replication at which it acted. We infected TRIM22 depleted or control MRC5t cells with IAV (PR8; MOI of 0.05 PFU/cell based on titres derived from MRC5t cells) in the presence of cycloheximide to inhibit viral protein synthesis and genome replication. Nuclei were isolated at 4 hpi and the relative levels of input vRNA were quantified by qRT-PCR. Western blotting for histone H3 and actin confirmed successful cell fractionation (Fig 6A). TRIM22 depletion did not significantly alter the nuclear accumulation of IAV genomes (Fig 6B), demonstrating that constitutive TRIM22 expression was not a barrier to viral genome entry into the nucleus of infected cells.

**FIG 6.**
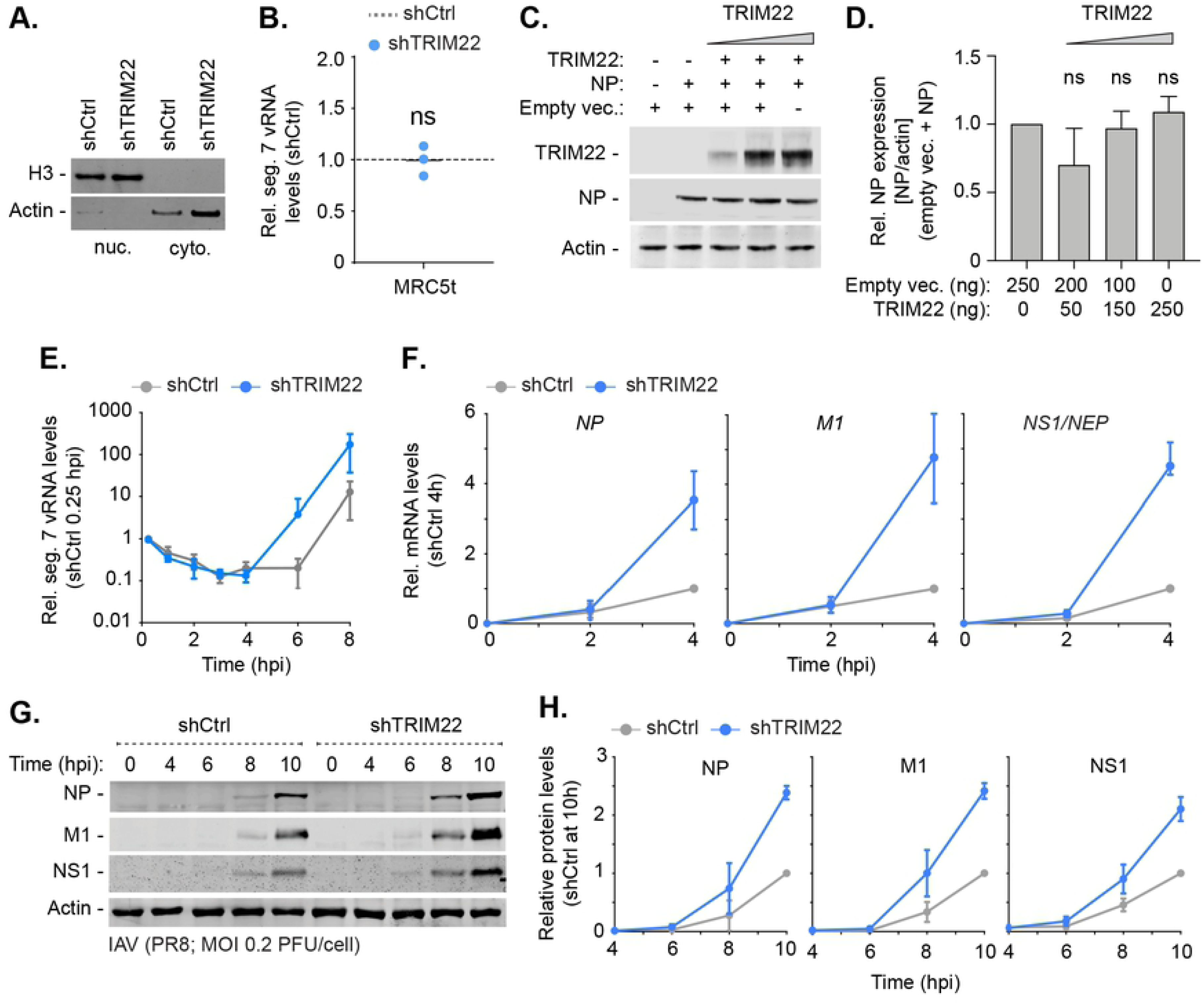
Constitutive TRIM22 expression restricts IAV transcription. (A-B) MRC5t shCtrl and shTRIM22 cells were infected with IAV (PR8) at a MOI of 0.05 PFU/cell (based on MDCK titres) in the presence of cycloheximide for 4 h prior to nuclear (nuc.) or cytosolic (cyto.) fractionation. (A) Western blot analysis showing histone H3 (nuc.) and actin (cyto.) expression profiles in fractionated lysates. (B) qRT-PCR quantitation of IAV PR8 seg. 7 vRNA levels in nuclear fraction lysates. Values normalized to shCtrl per experiment (dotted line). All data points shown; line: mean. One-sample two-tailed t test (hypothetical mean of 1; ns, not significant). (C, D) HEK 293T cells were transfected with increasing amounts of myc-tagged TRIM22 expression plasmid, 150 ng of NP (WSN) expression plasmid, and varying concentrations of empty vector control to balance DNA content for 24 h. (C) Western blot analysis of WCLs showing TRIM22 and NP expression levels. Actin is shown as a loading control. (D) Quantitation of NP expression protein levels (as in C). Values normalized to actin and expressed relative to NP in the absence of TRIM22. n=3, means and SD shown. (E-H) MRC5t shCtrl and shTRIM22 cells were infected with IAV (PR8) at a MOI of 0.05 (E) or 0.2 (F-H) PFU/cell (based on MRC5t titres) and harvested at the indicated time points. (E) Quantification of IAV vRNA seg. 7 levels by qRT-PCR. Values normalized to infected shCtrl samples at 0.25 hpi. n=3, means and SD shown. (F) qRT-PCR quantitation of IAV *NP*, *M1*, and *NS1/NEP* mRNA levels. Values normalized to infected shCtrl samples at 4 hpi. n=3, means and SD shown. (G) Western blots of infected WCLs showing viral (NP, M1, and NS1) protein expression levels. Actin is shown as a loading control. (H) Quantitation of viral protein expression levels (as in G). Values normalized to actin and expressed relative to levels in infected shCtrl cells at 10 hpi. n=3, means and SD shown.

We next asked whether TRIM22 might affect the stability of incoming viral genomes via degradation of NP, as transient transfection studies in HEK 293T cells have shown TRIM22 to target NP for ubiquitination and proteasome-mediated degradation [55, 61]. However, we were unable to detect any alteration of NP accumulation caused by TRIM22 in HEK 293T cells using an equivalent assay (Fig 6C, D). Nor were we able to detect any difference in genome stability between TRIM22 depleted or control MRC5t cells within the first four hours of IAV infection, prior to the onset of viral genome replication (Fig 6B and Fig 6E, 0 to 4 hpi). Thus, endogenous levels of constitutive TRIM22 expression can restrict IAV replication without influencing the stability of incoming viral ribonucleoproteins (vRNPs). However, following the onset of *de novo* vRNA synthesis higher levels of vRNA accumulated in TRIM22 depleted cells compared to control cells (Fig 6E, 4 to 8 hpi). These data suggested that endogenous TRIM22 affects viral genome replication. In order to replicate, IAV genomes must be encapsidated by newly synthesized viral proteins [35]. We therefore used qRT-PCR and western blotting to measure the transcription and expression profiles of the viral NP, M1 and NS1 genes, each encoded by independent genome segments, over a time course of infection. TRIM22 depletion increased transcription of all three genes (Fig 6F), correlating in each case with an increase in viral protein synthesis (Fig 6G, H). While transcription and replication of the IAV genome are intimately linked, the differences in mRNA levels were detectable prior to differences in *de novo* vRNA synthesis under equivalent infection conditions (Fig 6E, F). Thus, TRIM22 provides intrinsic immunity to IAV infection by suppressing the onset of viral transcription to restrict, either directly or indirectly, the initiation of viral genome replication. Collectively, these data highlight that constitutively expressed TRIM22 inhibits IAV replication early in the infectious cycle (2 to 4 hpi; Fig 6F) prior to the accumulation of viral immuno-evasion genes (for example NS1, Fig 6G, H; [13, 14]).

### Human lung tissue is enriched for constitutive ISG expression

Having identified TRIM22 to be a constitutively expressed ISG product within the respiratory tract that confers protection to IAV infection (Fig 1, 3, 5), we next examined whether other ISGs were constitutively expressed to high levels in the lung relative to other mucosal (gastrointestinal tract; esophagus, colon, and small intestine) or non-mucosal (liver, skin, and kidney) tissues. Using GTEx project (https://gtexportal.org/home/; [64]) RNA-seq data obtained from human tissue biopsies, we examined the transcript expression profiles of 200 ISGs previously identified to be upregulated in response to universal IFN stimulation in primary cell culture (≥ 8-fold change; [67]) (Fig 7, S2A Table). This analysis showed that individual tissues expressed distinct profiles of ISG transcript expression (Fig 7A, ranked by lung expression), with TRIM22 amongst the top 50 most abundantly expressed ISGs in the lung (Fig 7B, ranked by lung expression). Significantly higher levels of median ISG transcript expression were observed in the lung relative to all other tissues examined (Fig 7C, 200 ISGs). Principle component analysis (PCA) and clustering demonstrated that individual ISGs were not equally expressed in all tissues (Fig 7D), with lung tissue sharing the highest degree of ISG profile similarity to that of the small intestine (Fig 7E). Transcript levels of type-I (α, β), -II (γ), and -III (λ) IFNs were either not detectible or extremely low (≤ 1.3 TPM; Fig 7F, G, S2B Table), indicating that ISG enrichment in these tissues occurred independently of high levels of constitutive IFN transcription. While a role for IFN in the tissue-dependent enrichment of specific ISGs cannot be ruled out [18, 68, 69], these data demonstrate the existence of tissue-specific profiles of constitutive ISG transcription and show that lung tissue is enriched in ISG transcripts relative to other tissues.

**FIG 7.**
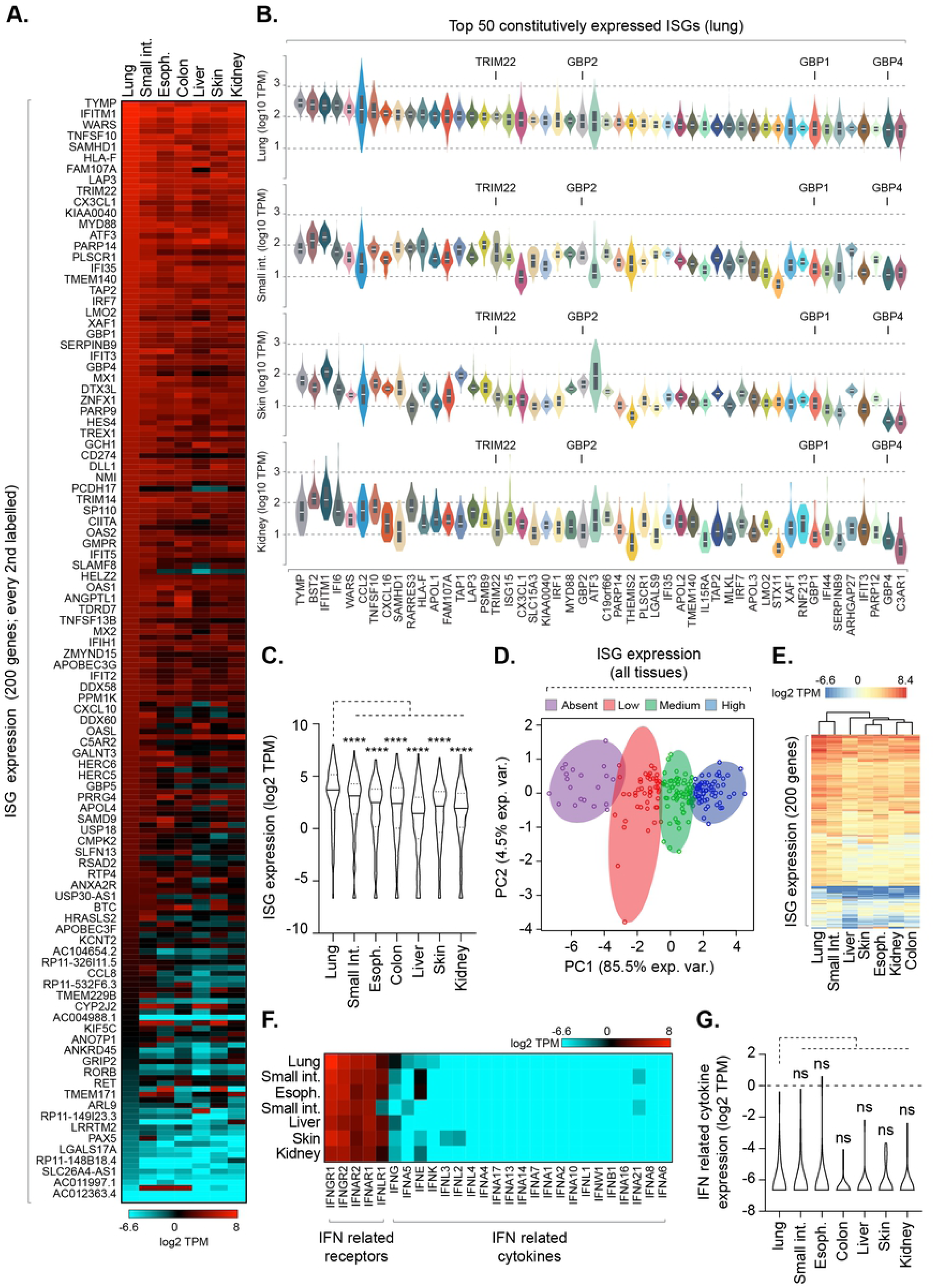
Lung tissue is enriched for constitutive ISG expression. (A) log2 median TPM expression values of 200 ISGs [67] across a range of human tissue biopsies (n); lung (n=427), small intestine (Int.; terminal ileum; n=137), esophagus (Espho.; mucosa; n=407), colon (sigmoid; n=233), liver (n=175), skin (suprapubic; n=387), and kidney (cortex; n=45). Every second gene labelled (full gene list in S2A Table). (B) Violin plots showing individual ISG expression profiles of the top 50 constitutively expressed lung ISGs and associated expression profiles in small intestine, skin, and kidney tissues. White line; median. Box; 5^th^ and 95^th^ percentile range. (C) Violin plot showing tissue expression profile of 200 ISGs across human tissues (as in A). Horizontal solid lines; median ISG expression per tissue. Horizontal dotted lines; 5^th^ and 95^th^ percentile range per tissue. (D) Principle component (PC) plot showing clustered ISG expression profiles across all tissues. (E) Heatmap showing clustered ISG transcript expression profiles between tissues. (F) log2 median TPM tissue expression values of IFN-related receptors and cytokines across human tissues (as in A; S2B Table). (G) Violin plot showing the median tissue expression values of IFN-related receptors and cytokines across human tissues (as in F). Horizontal solid lines; median. Horizontal dotted lines; 5^th^ and 95^th^ percentile range. Paired one-way ANOVA (Friedman multiple comparison test); **** *P* < 0.0001; ns, not significant. RNA-seq data adapted under creative commons licence from GTEx portal (https://gtexportal.org/home/; [64]).

Notably, many well established antiviral ISGs (BST2, IFITM1, and SAMHD1) and IAV associated host restriction factors, including the GBP (guanylate-binding protein) family, were observed to have high levels of constitutive ISG transcript expression in the lung (Fig 7B) [70, 71]. These data suggest that enriched levels of pre-existing ISG expression in the lung may combine to confer enhanced antiviral protection against respiratory airway infection immediately upon pathogen entry into susceptible host cells.

### The disruption of intracellular immune networks in transformed cells increases permissivity to IAV replication

Since the constitutive expression of TRIM22 is lost in many transformed cell lines (Fig 2A-D, S3 Fig), we hypothesized that other constitutively expressed ISGs might also be downregulated. Using cell-line RNA-seq data sets obtained from HPA (https://www.proteinatlas.org; [62, 63]), we compared the ISG transcript expression profile of non-treated hTERT-immortalized human bronchial epithelial cells (HBEC3-KT; HBEC3) to that of three widely used transformed cell lines (A549, HEK 293, and HeLa). Out of the 200 ISGs previously examined (Fig 7A), 178 ISGs were identified in RNA-seq cell line data sets (S3A Table) with 87 ISGs having values ≥ 5 TPM (Fig 8A; median 3.5 TPM per gene across all cell lines). Relative to HBEC3 cells, many of these ISGs were downregulated in transformed cells in an ISG-specific and cell-line dependent manner (Fig 8B; HBEC3 ≥ 5-fold change). Out of the 34 ISGs identified to be differentially downregulated in transformed cells (S3B Table), several were downregulated in all three (defined as core) or in two (defined as shared) transformed cell lines (Fig 8B, C). Importantly, the profile of ISG expression in unstimulated HBEC3 cells for this subset of ISGs was similar to that observed in lung tissue (Fig 8B; 34 ISGs); although the overall ISG expression profile significantly varied between lung tissue and all cell lines examined (S4 Fig; 178 ISGs). Network analysis using STRING (https://string-db.org; [72]) demonstrated that many of these downregulated ISGs were connected in the immune system network (17 of 34 genes; Fig 8C, D), and as a gene set to show pathway enrichment for defence response to virus and IAV infection (Fig 8D). Collectively, these data demonstrate that transformed cells display lineage-specific patterns of constitutive ISG expression, with a significant number of ISGs being downregulated relative to HBEC3 cells or lung tissue.

**FIG 8.**
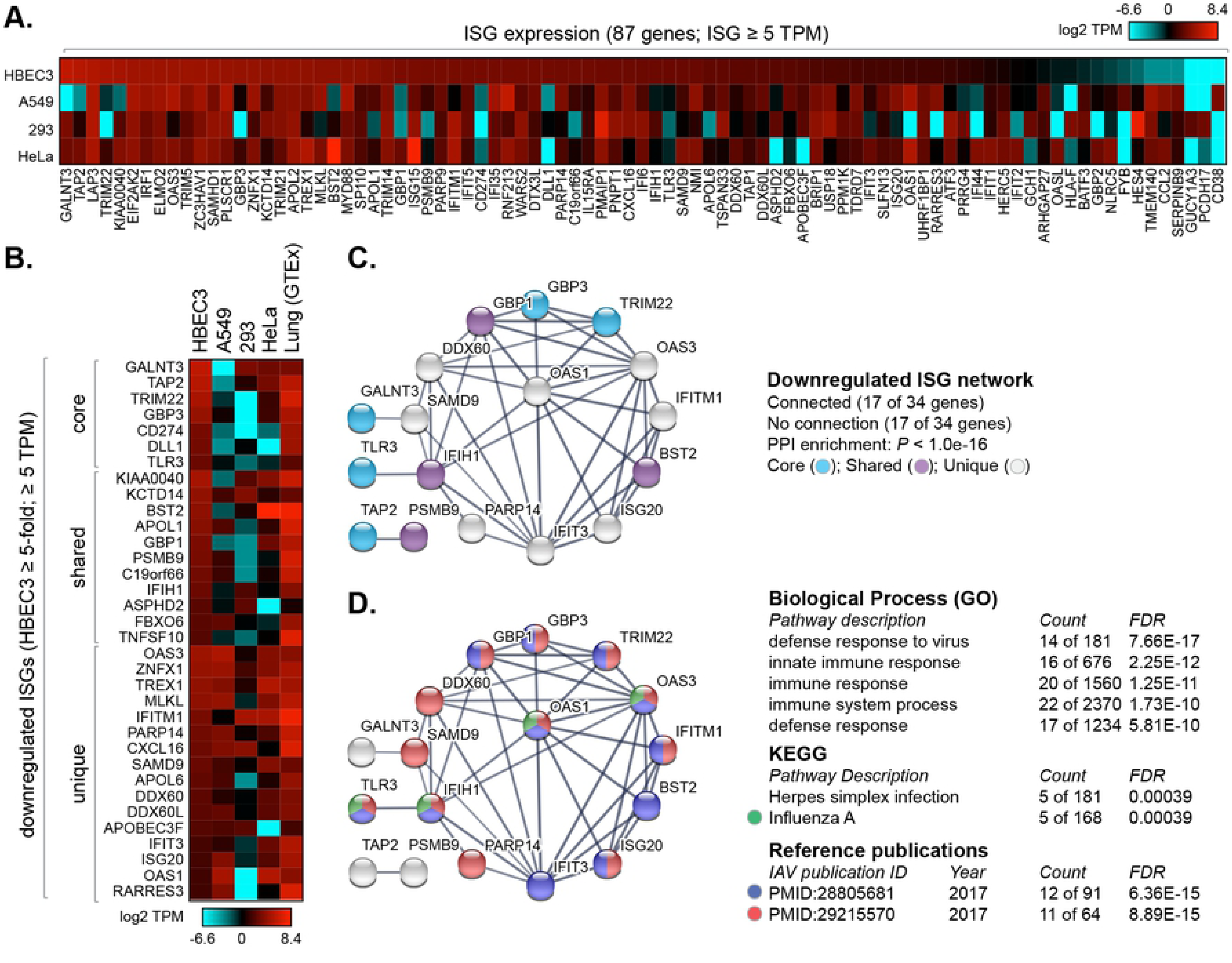
Virally-transformed cells have downregulated levels of constitutive ISG expression. (A) log2 TPM expression values of constitutive expressed ISGs (≥ 5 TPM per gene; 87 of 178 genes, S3A Table) in non-treated HBEC3, A549, HEK 293 (293), and HeLa cells. (B) Differentially downregulated ISGs (HBEC3 ≥ 5-fold change; as in A) plotted against the equivalent log2 median TPM ISG tissue expression values obtained from human lung tissue biopsies (n=427) (S3B Table). Downregulated ISGs common to three transformed cell lines; core: common to two transformed cell lines; shared: unique to one transformed cell line; unique. (D/E) High-confidence (> 0.7) STRING (https://string-db.org; [72]) protein-protein interaction (PPI) network of identified downregulated ISGs (as in B). Core; blue circles: shared; purple circles: unique; grey circles. Network PPI enrichment: *P* < 1.0e-16. Ranked biological process (GO), KEGG pathways (green circles in D), and reference publications (red and blue circles in D; [70, 71]) with associated counts in gene sets (count) and FDR (false discovery rate) values shown. Cell line and tissue RNA-seq data adapted under creative commons licence from HPA and GTEx portal, respectively [62–64].

Having identified a subset of constitutively expressed ISGs known to restrict IAV to be downregulated in transformed cells (Fig 8), we extended our analysis to determine whether other immune system-related genes were downregulated relative to HBEC3 cells (HBEC3 ≥ 5-fold change; blue circles in Fig 9A; S4A Table). Gene Ontology (GO) analysis identified that a significant percentage of differentially downregulated genes in A549 (18.09%), HEK 293 (17.39%), and HeLa (16.46%) cells map to the immune system (S4B, C Tables). Out of the 174 unique immune genes identified to be downregulated, 95 (54.6%) were common to at least two transformed cell lines (Fig 9B, core + shared; S4D Table). STRING analysis identified a significant degree of network connectivity between these downregulated immune genes (141 of 174 genes; Fig 9C, S5 Fig), with common (core + shared) immune genes located across the entire network. We conclude that transformed cells share common networks of immune system disruption which arise through lineage-specific patterns of immune gene downregulation. Notably, this gene network was also enriched for host factors known to restrict IAV (Fig 9C; KEGG pathway [<u>hsa05164</u>]), suggesting that transformed cells are deficient in multiple host factors known to contribute to the intracellular restriction of IAV.

**FIG 9.**
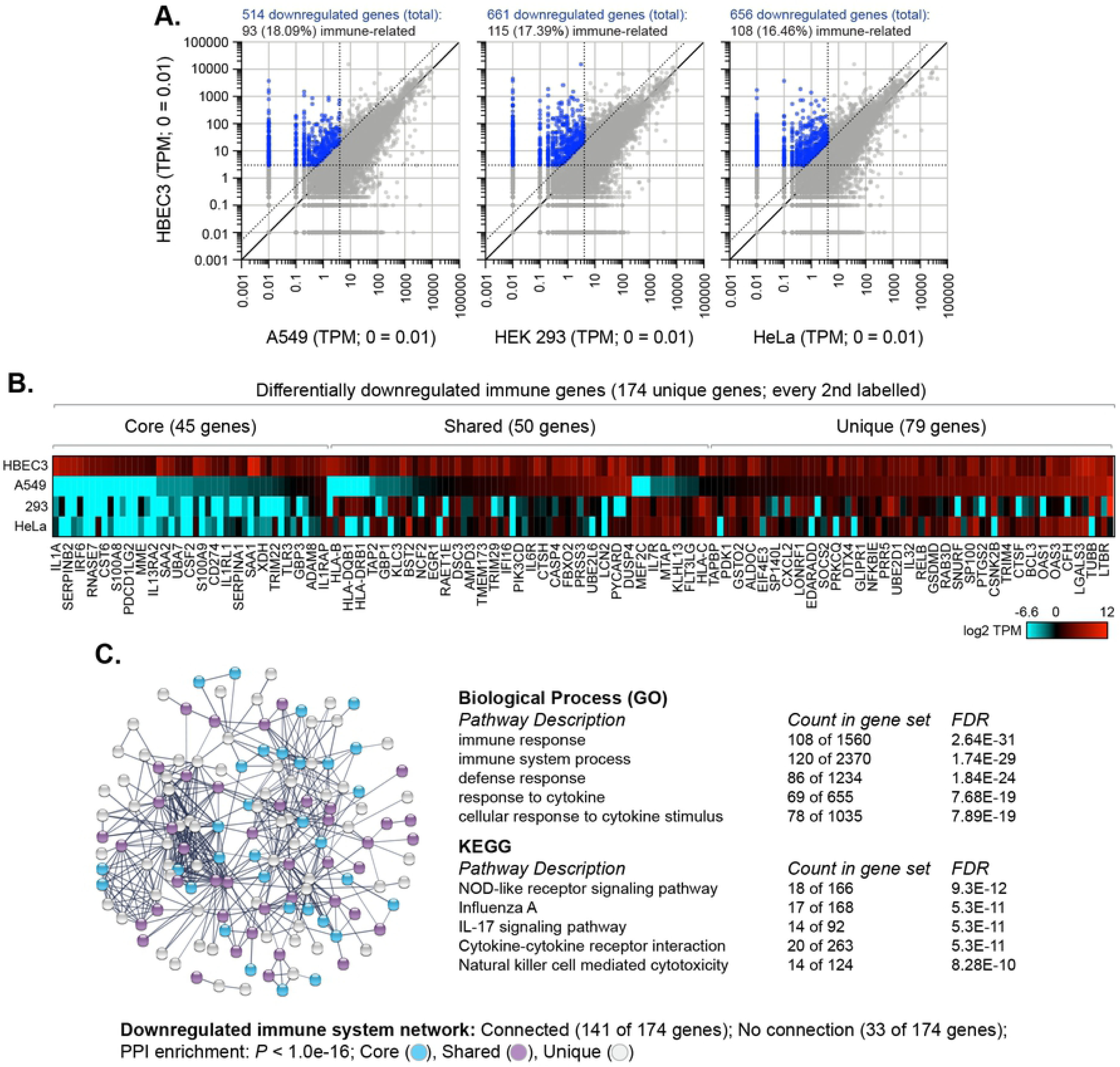
Transformed cells are deficient in the constitutive expression of multiple immune regulators. (A) Scatter plots highlighting downregulated genes (blue circles) identified between non-treated HBEC3 cells (≥ 3 TPM; median HBEC3 TPM per gene, horizontal dotted line) and A549 (median 3 TPM per gene), HEK 293 (293; median 3 TPM per gene), or HeLa cells (median 4 TPM per gene; ≤ 4 TPM, vertical dotted line; HBEC3 ≥ 5-fold change, diagonal dotted line; S4A Table). Differentially downregulated genes were mapped and used for pathway analysis using Reactome (https://reactome.org; Table S4B). The number of downregulated genes and percentage mapped to the immune system (% immune-related; Table S4C) is shown. (B) log2 TPM gene expression profiles of downregulated immune system genes (as in A; 174 unique genes identified, every second labelled). Downregulated genes common to three transformed cell lines; core: common to two transformed cell lines; shared: unique to one transformed cell line; unique (as highlighted; S4D Table). (C) High-confidence (> 0.7) STRING protein-protein interaction (PPI) network of identified downregulated immune system genes (as in B). Core; blue circles: shared; purple circles: unique; grey circles. Network PPI enrichment: *P* < 1.0e-16. Ranked biological process (GO) and KEGG pathways with associated counts in gene sets (count) and FDR (false discovery rate) values shown. An enlarged annotated map is presented in S5 Fig. Cell line RNA-seq data adapted under creative commons licence from HPA [62, 63].

To investigate this observation further, we curated an extended IAV KEGG network which included recently identified host factors that influence IAV restriction (Fig 10A, S5A Table; [70, 71]). Out of the 184 genes analyzed, 39 (21.2%) were identified to be significantly downregulated in transformed cells relative to HBEC3 cells (≥ 5-fold change; Fig 10B, C, S5B Table) or lung tissue. Consistent with both ISG and immune system profiling (Fig 8C, 9B, respectively), many of these genes were downregulated in two or more transformed cell lines and showed significant network connectivity (29 of 39 genes; Fig 10D). The expression profiles of a subset of these proteins were tested in unstimulated cells by western blotting, which confirmed that UBA7, TRIM22 (positive control), IFITM1, GBP1, IFIH1 and TLR3 were expressed to significantly lower levels in A549 cells relative to HBEC3 cells (Fig 10E, F). In order to determine if the disruption of this immune system network influenced IAV replication, we compared the relative plaque titre of IAV in HBEC3 and A549 cells to that of MDCK cells. Similar to diploid lung fibroblasts (Fig 2G, H), human bronchial epithelial cells were highly restrictive to the initiation of IAV plaque formation relative to MDCK cells (≥ 70-fold) or A549 cells (≥ 30-fold) (Fig 9G, H). Ruxolitinib inhibition of JAK-STAT signalling did not influence the initiation of IAV plaque formation in any of the cell-types examined (Fig 9I), although a significant increase in plaque diameter could be observed in each cell-type (Fig 9J). Thus, pharmacological inhibition of cytokine-mediated innate immune defences enhances virus propagation and spread, but not the initiation of viral replication leading to plaque formation [11, 66]. We conclude that the constitutive expression of IAV immuno-regulatory genes in the context of non-transformed respiratory cells confers a significant pre-existing immune barrier to IAV infection prior to the induction of pathogen-induced cytokine-mediated innate immune defences. Importantly, this intrinsic barrier is compromised in many transformed cell lines currently being used for IAV immunobiology research.

**FIG 10.**
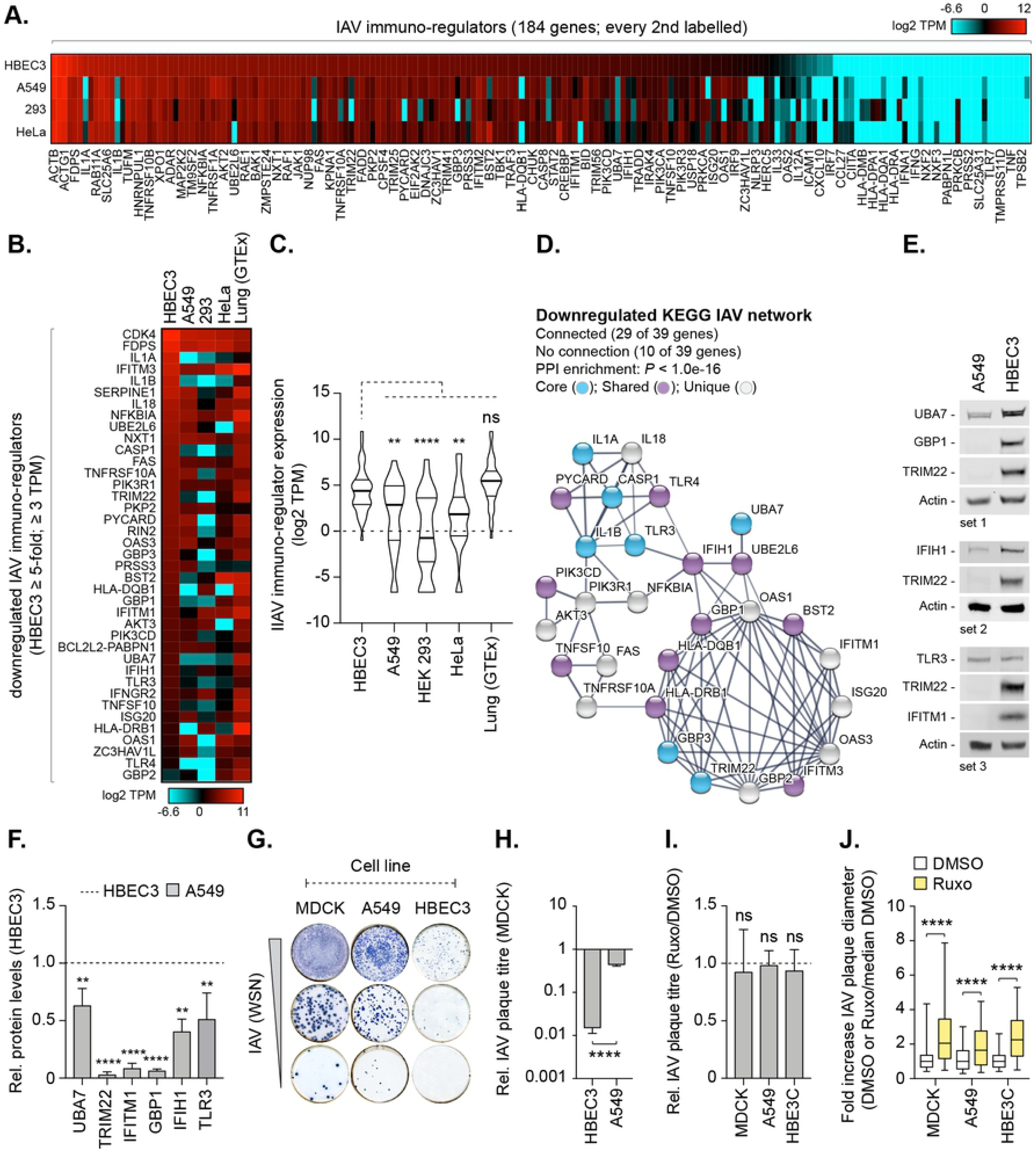
Transformed cells are permissive to IAV replication independently of pathogen-induced cytokine-mediated innate immune defences. (A) log2 TPM expression values of 184 IAV immuno-regulatory genes (extended IAV KEGG network (<u>hsa05164</u>); [70]; S5A Table) in non-treated HBEC3, A549, HEK 293 (293), and HeLa cells. Every second gene labelled. (B) Differentially downregulated IAV immuno-regulatory IAV genes (HBEC3 ≥ 5-fold change, ≥ 3 TPM per gene) plotted against the equivalent log2 median TPM gene expression values obtained from human lung tissue biopsies (n=427) (S5B Table). (A, B) Cell line and tissue RNA-seq data adapted under creative commons licence from HPA and GTEx portal, respectively [62–64]. (C) Violin plot showing log2 TPM expression values of downregulated IAV immuno-regulatory genes (as in B). Horizontal solid lines; median gene expression. Horizontal dotted lines; 5^th^ and 95^th^ percentile range. Paired one-way ANOVA (Friedman multiple comparison test); ** *P* < 0.01; **** *P* < 0.0001; ns, not significant. (D) High-confidence (> 0.7) STRING protein-protein interaction (PPI) network of identified downregulated KEGG associated IAV genes (as in B). Core; blue circles: shared; purple circles: unique; grey circles. Network PPI enrichment: *P* < 1.0e-16. (E) Western blots of non-treated HBEC3 and A549 WCLs showing UBA7, GBP1, IFIH1, TLR3, IFITM1, and TRIM22 (+ve control) protein expression levels. Actin is shown as a loading control. (F) Quantitation of protein expression levels (as in E). Values normalized to actin and expressed relative to levels in HBEC3 cells. n≥3, means and SD shown. One-sample two-tailed t test (hypothetical mean of 1; ** *P* < 0.005, **** *P* < 0.0001). (G) Representative immunocytochemistry images of IAV plaque formation (NP staining) in MDCK, A549, and HBEC3 cells infected with equivalent serial dilutions of IAV (WSN). (H) Quantitation of plaque numbers in each cell line expressed relative (rel.) to MDCK cells (rel. plaque titre); n≥3, means and SD shown. Mann-Whitney U-test; **** *P* < 0.0001. (I) MDCK, A549, and HBEC3 cells were pre-treated for 1 h in the presence of Ruxolitinib (Ruxo; 5 µM) or DMSO (carrier control) prior to infection with serial dilutions of IAV (WSN; in the presence of drug or carrier control). Quantitation of rel. plaque titre (plaque titre with Ruxo / plaque titre with DMSO) for each cell line is shown. n≥3, means and SD shown. One-sample two-tailed t test (hypothetical mean of 1; ns, not significant). (J) Quantitation of the fold increase in IAV plaque diameter in Ruxo or DMSO-treated infected cell monolayers (as in I). Values normalized to the median DMSO plaque diameter in each cell line (DMSO or Ruxo/median DMSO). n≥3, means and SD shown. Mann-Whitney U-test; **** *P* < 0.0001.

## Discussion

Recent single-cell transcriptomic studies have identified individual cells to be differentially permissive to IAV infection [24, 25]. These observations suggest that pre-existing (intrinsic) patterns of cellular gene expression within (or between) specific cell-types may differentially influence the outcome of IAV infection. However, biological evidence to support the importance of such cell-type specific patterns of host gene expression in the intracellular restriction of IAV has remained lacking. Consequently, the concept of intrinsic immunity has yet to be firmly accepted within the IAV field or wider respiratory virus research community. Here, we demonstrate that non-transformed human lung cells possess patterns of constitutive antiviral gene expression that differ markedly from the transformed cell culture model systems that have been widely used for respiratory virus research over many decades. We show that these differential patterns of constitutive antiviral gene expression can directly influence the outcome of IAV infection, independently of pathogen-induced cytokine-mediated innate immune defences. Thus, we have identified a biologically important and previously overlooked role for intrinsic immunity in the regulation of IAV infection; findings relevant to the intracellular immune regulation of many respiratory pathogens.

Our initial hypothesis that cells in the respiratory mucosa might differentially express antiviral proteins at higher levels than in cells at less exposed locations led to the identification of TRIM22 to be amongst the most abundantly expressed TRIM proteins in lung tissue and non-transformed cells of lung origin (Fig 1, 2A, S1, S6 Fig). This high level of constitutive TRIM22 expression contrasts with many previous studies, which have reported TRIM22 to be strongly upregulated as an effector ISG in primary lymphocytes and transformed cell lines in response to virus infection or immune stimulation (Fig 2, S3 Fig; [54, 55]). We note that TRIM22 (formerly known as Staf50) has been shown to be upregulated by p53 and its expression has been found to correlate with cell differentiation and proliferation status [54, 73]. Thus, the downregulation of constitutive TRIM22 expression in many virally-transformed cells may occur as a direct result of p53 inactivation by viral proteins to sustain cellular proliferation. However, it is evident that many transformed and carcinoma cell lines have variable gene copy numbers and an extensive array of single nucleotide polymorphisms (SNPs) that can directly influence cellular protein expression profiles, protein functionality, and immune competence [74–76]. Indeed, analysis of Cancer Cell Line Encyclopaedia (CCLE; [75]) records demonstrate that TRIM22 transcript levels and copy number are downregulated in many lung carcinoma cell-types (https://portals.broadinstitute.org/ccle/page?gene=TRIM22). Such issues of genetic variance raise concerns over the suitability of using carcinoma cells for virus-related immunity studies. For example, it is becoming increasingly evident that many cancers downregulate multiple immune regulators to minimize immune clearance and carry unique epigenetic signatures that influence gene transcriptional regulation and proliferation [74, 75, 77, 78]. Consequently, the utilization of such genetically variable populations of cells for *in vitro* experimentation is likely to have a significant bearing on viral intracellular immune regulation due to lineage-specific patterns of immune gene regulation acquired through transformation.

We demonstrate that multiple immune regulators, known to influence the replication of a wide variety of viral pathogens, are downregulated in transformed cell lines widely used for respiratory virus research (Fig 8-10). The loss or downregulation of these constitutively expressed host factors correlates strongly with enhanced permissivity of these cell-types to the initiation of IAV replication leading to plaque formation (Fig 2G, 10H). Many of these immune genes, although downregulated in a lineage-specific manner (Fig 9), share common networks of immune-system regulation known to influence IAV replication (Fig 8C, 9C, 10D). These observations may account for much of the gene-specific variability, but interrelated pathway connectedness, observed between genome-wide RNA interference screens that utilized carcinoma model systems to identify host factors that influence IAV replication [79–81]. Collectively, our data highlight the importance of utilizing more physiologically relevant cell culture model systems to improve experimental reproducibility between independent groups and research fields.

Using a cell culture system that retained the constitutive expression of TRIM22 observed at the natural site of infection (Fig 1, 2), we corroborate previous reports that TRIM22 acts as a restriction factor to inhibit IAV replication (Fig 3F-H; [55, 61]). Importantly, we show that constitutive expression of TRIM22 is sufficient to restrict the initiating cycle of both human and avian IAV replication from the outset of infection by inhibiting the efficient onset of viral transcription (Fig 5, 6). We show that pharmacological inhibition of cytokine-mediated JAK-STAT signalling did not reduce the ability of endogenous TRIM22 to restrict IAV infection (Fig 4; [66]), demonstrating the TRIM22 can work independently of the IFN pathway. Thus, high levels of constitutive TRIM22 expression can confer immediate protection to airway infection, thereby reducing the need to prematurely activate potentially harmful pro-inflammatory innate immune defences [2, 33, 34].

Our work exemplifies two important features of intrinsic immunity. Firstly, we identify TRIM22 to be an intrinsically-expressed ISG, similar to PML (TRIM19) and TRIM5α [19, 46, 48, 49], which can be further upregulated in response to cytokine signalling in a manner dependent on the pre-existing basal levels of endogenous expression in a given cell-type (Fig 1F-H, 2A, B, S3 Fig). Secondly, TRIM22 demonstrates that intrinsic immune defences can be upregulated in a tissue-specific manner. Like TRIM32 and TRIM41 [21, 22], TRIM22 can restrict the initiation of IAV infection (Fig 5, 6), but unlike these TRIM proteins TRIM22 is upregulated in a tissue-specific manner, being enriched within the lung relative to other tissues (Fig 1A, B, 7A, B). Collectively, these observations point to a series of distinct ways in which constitutive levels of immune gene expression can influence the outcome of IAV replication independently of pathogen-induced host defences. Further investigation is warranted to determine the accumulative and strain-dependent effects of such intrinsic barriers to IAV infection.

We found that constitutively-expressed TRIM22 could restrict the replication of multiple human (H1N1, including WSN) and avian (H3N2 and H7N1) strains of IAV (Fig 5). These data contrast with recent studies in transformed cells [61], which lack constitutive TRIM22 expression (Fig 2A, S3; [55, 61]), that reported the WSN strain to be resistant to TRIM22 mediated restriction. In these studies, TRIM22 was found to target IAV NP for ubiquitination and proteasome-dependent degradation [55, 61], with WSN NP being resistant to ubiquitination due to the substitution of lysine acceptor residues for arginines [61]. In contrast, we found that constitutively-expressed TRIM22 was effective against the WSN strain (Fig 3F-H) independently of detectible NP degradation, either alone or in the context of incoming vRNPs (Fig 6C-E). While we cannot discount a role for ubiquitination in the TRIM22 mediated restriction of IAV, we show endogenous levels of TRIM22 are sufficient to restrict *de novo* NP expression by inhibiting the onset of viral transcription (Fig 5, 6).

These differences may reflect cell-type (transformed *vs* non-transformed cells) or expression level (ectopic *vs* endogenous) dependent differences in TRIM22 restriction of IAV and suggest that TRIM22 may adopt multiple approaches to restrict IAV infection. For example, endogenous levels of TRIM22 may sterically hinder the onset of viral transcription independently of NP degradation when in complex with vRNPs, but upon saturation under high genome loads to target *de novo* synthesized free pools of NP for ubiquitination leading to its proteasome-degradation. Such differences in substrate-targeting could result in a switch in intracellular immune ‘status’ from intrinsic to innate defences, induced by the onset of viral replication or sensing of PAMPs in a strain-dependent manner. Such a mechanism is not unprecedented, as we have recently shown that PML (TRIM19) plays spatiotemporally distinct roles in the regulation of intrinsic and innate immune defences to HSV-1 infection [19, 82]. Further biochemical investigation will be required to determine whether TRIM22 has differential modes of substrate-targeting dependent on the kinetics of infection or cellular immune status.

A surprising discovery in our study was the identification of enriched levels of constitutive ISG expression in human lung tissue relative to that of other mucosal and non-mucosal tissues (Fig 7). Our analysis suggests that human lung tissue could confer heightened levels of pre-existing immune protection against multiple respiratory viruses immediately upon pathogen entry. Importantly, this was not due to the elevated expression of all ISGs, but rather tissue-specific profiles of individual ISG expression (Fig 7A-E, S2A Table). These data demonstrate that human tissues confer distinct profiles of ISG expression in a tissue-dependent manner which may confer enhanced protection at exposed surfaces.

While it remains to be determined how such distinct patterns of ISG expression occur, some plausible explanations include: (i) the presence of commensal microbiota or natural turnover of cells, stimulating low levels of PRR activation and IFN secretion through the release of PAMPs or DAMPs, respectively [18, 83]; (ii) differential patterns of cytokine secretion between tissues, including low basal levels of ‘tonic’ IFN signalling to maintain immune-readiness and fitness [68, 69, 84, 85]; (iii) tissue-specific patterns of transcriptional regulation occurring through cellular differentiation, which may be further influenced by inherited genetic traits or epigenetic status [85–88]. Importantly, these explanations are not mutually exclusive, which may account for the variance in individual ISG expression profiles observed between human tissue samples (Fig 7B). Further work is required to determine how such tissue-specific signatures of pre-existing antiviral gene expression influence the initiation and outcome of respiratory virus infection.

In conclusion, we identify pre-existing tissue-specific and cell-type dependent patterns of constitutive immune gene expression which confer a significant intracellular immune barrier to IAV replication from the outset of infection and independently of pathogen-induced cytokine-mediated innate immune defences. These intrinsic barrier defences are downregulated in many transformed cell lines currently used for respiratory virus research, which share common networks of immune system disruption relevant to the immune regulation of many respiratory pathogens.

## Materials and Methods

### Antibodies

Polyclonal antibodies were used to detect TRIM22 (Sigma-Aldrich; HPA003575), Mx1 (Santa Cruz; sc-50509), GBP1 (Proteintech; 15303-1-AP), IFIH1 (Proteintech; 21775-1-AP), TLR3 (Proteintech; 17766-1-AP), histone H3K27ac (AbCam; ab4729), and actin (Sigma-Aldrich; A5060). IAV hybridoma antisera were used to detect NP, M1, and NS1, as previously described [89]. Monoclonal antibodies were used to detect UBA7 (AbCam; ab133499), Actin (DSHB; 224-236-1), IFITM1 (Proteinech; 60074-1g), and IAV NP (AbCam; ab20343). Secondary antibodies were Alexa 488 and 555 donkey anti-mouse and - rabbit (Invitrogen; A21202, A21206, and A31572), DyLight 680- or 800-conjugated anti-rabbit (Thermo Fisher Scientific; 35568 and SA5-35571), and peroxidase conjugated anti-mouse (Sigma-Aldrich; A4416).

### Animals and ethics

No animals were directly subjected to experimentation as part of this scientific study. All animal tissues were obtained from material produced in previously described experiments [65] with permission from Public Health England (PHE). Procedures associated with this earlier study were approved by the PHE Ethical Review Committee (Porton Down, UK) and authorized under UK Home Office project licence 30/3083.

### Quantitative Histopathology of cynomolgus macaque tissue sections

Formalin fixed and paraffin embedded tissue samples were processed for haematoxylin and TRIM22 immunohistochemistry (IHC) staining, as previously described [65]. Tissue sections were independently assessed for TRIM22 expression by a qualified pathologist. Automated quantitation of TRIM22 expression levels in stained tissue sections was performed using whole-slide scans and Image-Pro Premier (Media Cybernetics), as previously described [90, 91].

### Quantitative Histopathology of human tissue sections

IHC data from human tissue samples was obtained from the Human Protein Atlas (HPA; http://www.proteinatlas.org; [62, 63]) under a Creative Commons Attribution-ShareAlike 3.0 International License. The original images consulted were TRIM22 Bronchus (http://www.proteinatlas.org/ENSG00000132274-TRIM22/tissue/bronchus#img) and TRIM22 Nasopharynx (http://www.proteinatlas.org/ENSG00000132274-TRIM22/tissue/nasopharynx#img).

### RNA-seq analysis of human cell lines and tissues

RNA-seq data for human cell lines (HBEC3-KT, A549, HEK 293, and HeLa) and human tissue biopsies (as indicated) were obtained from Human Protein Atlas (HPA; http://www.proteinatlas.org, version 18.1; [62, 63]) under a Creative Commons Attribution-ShareAlike 3.0 International License or Genotype-Tissue Expression (GTEx; https://gtexportal.org/home/, V7; [64]) project (as stated) supported by the Common Fund of the Office of Director of the National Institutes of Health, and by NCI, NHGRI, NHLBI, NIDA, NIMH, and NINDS. Principle Component Analysis (PCA) cluster plots were generated using kmeans and clusplot packages and prcomp function in R (https://www.r-project.org). Heatmaps were generated using Prism 8 (GraphPad) or pheatmap (v1.0.12) and cluster (v2.0.7-1) package in R. Network analysis was conducted using STRING (https://string-db.org; [72]).

### Cells, viruses, and drugs

Primary human foetal lung fibroblast (MRC5) cells were purchased from the European Collection of Authenticated Cell Cultures (ECACC; 05072101). MRC5t cells are immortalized MRC5 cells expressing the catalytic subunit of human telomerase (hTERT), and were generated as previously described [92]. MRC5 and MRC5t cells were cultured in Dulbecco’s Modified Eagle Medium (DMEM; Life Technologies; 41966) supplemented with 10 % foetal bovine serum (FBS; Life Technologies; 10270), 100 U/ml of penicillin and 100 μg/ml of streptomycin (P/S; Life Technologies; 15140-122), and 1× non-essential amino acids (NEAA; Life Technologies 11140-035). MRC5t cells were supplemented with 5 μg/mL hygromycin B (Thermo Fisher Scientific; 10687010) to maintain hTERT expression. MRC5t cells were transduced with lentiviruses to express short hairpin (sh) RNAs based on the 19-mer sequences; non-targeting control (shCtrl; 5’-TTATCGCGCATATCACGCG-3’) or TRIM22 targeting (shTRIM22 clone B7 [3’ UTR]; 5’-TATTGGTGTTCAAGACTAT-3’, clone B8; 5’-CTGTACGCACCTGCACATT-3’, clone B9; 5’-GTGTCTTCGGCTGCCAATA-3’), as previously described [46]. Pooled, stably transduced cells were maintained in growth media supplemented with 0.5 µg/ml puromycin (Sigma-Aldrich; P8833). Primary human bronchial epithelial (HBEp) cells were purchased from Sigma-Aldrich (502-05a). hTERT and CDK4 immortalized human bronchial epithelial cells (HBEC3-KT) were purchased the Hamon Center for Therapeutic Oncology Research (UT Southwestern Medical Center; [93]). Cells were cultured according to supplier guidelines. Madin Darby Canine Kidney (MDCK; a gift from Ben Hale University of Zurich), human lung adenocarcinoma epithelial (A549; PHE Culture Collections, 86012804), human embryonic kidney (HEK 293T; a gift from Roger Everett MRC-UoG CVR) and human cervical carcinoma (HeLa [a gift from Juergen Hass University of Edinburgh] or HEp2 [a gift from Roger Everett MRC-UoG CVR]) cells were cultured in DMEM with 10 % FBS and P/S. All cells were maintained at 37°C in 5 % CO_2_. IAV strains A/WSN/1933(H1N1) (WSN), A/Puerto Rico/8/1934(H1N1) (PR8), A/Udorn/307/1972(H3N2) (Udorn), and A/California/04/2009(H1N1) (Cal) were propagated in MDCK cells. A/Duck/Singapore/5/1997(H5N3) (Duck H5N3) and A/Chicken/Italy/1067/1999(H7N1) (Chicken H7N1) were propagated in embryonated chicken eggs. WSN titres were calculated by immunocytochemistry (ICC) plaque assay, as described below. PR8, Udorn, Duck H5H3 and Chicken H7N1 titres were calculated based on fluorescence forming units (FFU), calculated from the proportion of NP-positive MRC5t cells detected at 8 hours post-infection (hpi) by immunofluorescence confocal microscopy. Cells were interferon stimulated by the addition of 100 IU/ml recombinant interferon-β (IFN-β; Merck, 407318) to the growth media for 24 h. Cycloheximide (CHX; Sigma-Aldrich, C-7698) was prepared in Milli-Q water and used at 10 μg/ml. Ruxolitinib (Ruxo; Selleckchem; S1378) was prepared in DMSO and used at the concentrations indicated.

### Plaque and virus yield assays

For plaque assays, cells were seeded at 2 × 10^5^ cells/well in 12-well dishes and incubated for a minimum of 16 h prior to manipulation. Cells were infected with serial dilutions of virus for 1 h at 37°C prior to overlay with conditioned growth medium supplemented with 1.2 % Avicel (Biopolymers; RC-591), 0.1 % sodium bicarbonate (Life Technologies; 25080-060), and 0.01 % DEAE Dextran (Sigma-Aldrich; D9885). Cell monolayers were processed for ICC staining at 24 to 72 hpi depending on virus replication kinetics (as previously described [94]) or stained with Giemsa stain (VWR; 35086). Relative plaque titre was calculated as the plaque titre of a virus stock under the indicated condition divided by its titre under a control condition. For virus yield assays, cells were infected with IAV (WSN) at the indicated multiplicity of infection (MOI) for 1 h at 37°C, washed twice with PBS, then overlaid with growth medium. Supernatants were collected at the indicated time points post-infection and the released virus was titred by plaque assay in MDCK cells. Plaque diameters were measured using an automated Celigo imaging cytometer (Nexcelom biosciences), as per the manufacturer’s instructions.

### Immunofluorescence confocal microscopy

1 × 10^5^ cells were seeded on 13 mm glass coverslips in 24-well dishes and incubated for a minimum of 16 h prior to manipulation. After treatment, cell monolayers were fixed, permeabilized, and immunostained at the indicated time points, as previously described [94]. Nuclei were stained with DAPI (Sigma-Aldrich, D9542). Coverslips were mounted on glass slides using Citiflour AF1 mounting medium (AgarScientific; R1320) and sealed with nail enamel. Samples were examined with a Zeiss LSM 880 or LSM 710 confocal microscope with 405, 488, 543, and 633-nm laser lines. Images were captured under a Plan-Apochromat 63×/1.4 oil immersion or Plan-Neofluar 20×/0.5 air objective lenses. The proportion of IAV NP antigen positive cells was calculated from a minimum of five wide field images, imaging more than 1000 cells per coverslip per condition. The proportion of NP positive cells was determined and the fold increase in NP positive cells between IAV infected shTRIM22 and shCtrl MRC5t cells calculated for each biological repeat.

### Western Blotting

Cells were seeded at 2 × 10^5^ cells/well in 12-well dishes and incubated for a minimum of 16 h prior to manipulation. Treated or infected cell monolayers were washed twice in PBS and whole cell lysates (WCLs) collected in Laemmli buffer. Proteins were resolved on NOVEX NU-PAGE (4-12%) Bis-Tris gels (Invitrogen; NP0322), transferred onto 0.22 µm nitrocellulose membranes (Amersham; 15249794), and probed by western blotting, as previously described [94]. Membranes were imaged using an Odyssey Infrared Imager (Li-Cor). Band intensities were quantified using Image Studio Software (Li-Cor).

### qRT-PCR

For viral or cellular mRNA quantitation, cells were seeded at 2 × 10^5^ cells/well in 12-well dishes and incubated for a minimum of 16 h prior to manipulation. Treated or infected cell monolayers were washed once in PBS prior to RNA extraction using an RNAeasy Plus Kit (Qiagen; 74134). mRNA was reverse transcribed (RT) using the TaqMan Reverse Transcription Reagents kit (Life Technologies; N8080234) with oligo (dT) primers. Samples were analysed in triplicate using the TaqMan Fast Universal PCR Master Mix (Life

Technologies, 4352042) and TaqMan *GAPDH* (4333764F), *TRIM22* (Hs01001179_m1), *Mx1* (Hs00895608_m1) or *ISG15* (Hs01921425_s1) specific primer-probe (FAM/MGB; Thermo Fisher Scientific) mixes or custom IAV (*NP*, *M1*, *NS1*/*NEP*) primer-probes mixes (S7 Table). The ΔΔCt method was used to normalize transcript levels to those of *GAPDH* mRNA. For vRNA analysis, cells were seeded at 4 × 10^5^ cells/well in 6-well plates and incubated for a minimum of 16 h prior to manipulation. Cells were infected either in the presence or absence of CHX. At the indicated time points, cell monolayers were washed in PBS, harvested by trypsinization, and cell pellets washed twice in ice cold PBS. Nuclear and cytoplasmic fractions were isolated using NucBuster (Novagen 71183-3). If appropriate, fractions were divided for both western blot and qRT-PCR analysis. For vRNA analysis, total RNA was isolated from nuclear pellets using an RNAeasy Plus Kit. An IAV segment 7 specific primer (5’-AGCCGAGATCGCACAGAGACTT-3’) was used for reverse transcription, as previously described [95]. Samples were analysed in triplicate using the M1 primer-probe mix relative to a synthetic segment 7 (M) vRNA reference standard. The segment 7 vRNA standard was produced as previously described [95]. Briefly, vRNA was extracted from infected MDCK cells using the QIAamp Viral RNA Mini kit (Qiagen, 52904) and extracted RNA was reverse transcribed using the Uni12 universal IAV segment primer (5’-AGCAAAAGCAGG-3’) and TaqMan Reverse Transcription Reagent kit. The cDNA was used as a template to amplify the IAV segment 7 ORF, incorporating a T7 promoter sequence that was used to generate synthetic segment 7 vRNA using the TranscriptAID T7 high yield transcription kit (Thermo Fisher Scientific; K0441). Synthetic vRNA was purified using an RNAeasy column and used as a reference standard for reverse transcription and qRT-PCR analysis.

### Plasmids and transfections

A cDNA encoding wild-type (WT) human TRIM22 (a gift form Professor Juergen Hass, University of Edinburgh) was inserted into pcDNA.3.1 (Invitrogen) in frame with a 5’ Myc-tag oligo to generate a pcDNA.Myc.TRIM22. Clones were verified by Sanger sequencing. The IAV (WSN) pcDNA-NP (WSN) expression plasmid has been described previously [96, 97]. All transfections were performed using Lipofectamine 2000 (Thermo Fisher Scientific; 11668). For NP stability assays, 1 x 10^5^ HEK 293T cells/well were seeded onto poly-lysine (Sigma; 7405) coated 24-well plates. 24 h post-seeding, cells were co-transfected with 150 ng of pcDNA-NP (WSN) and 0, 100, 200, or 250 ng of pcDNA.Myc.TRIM22. Input levels of DNA were equalized by the inclusion of pcDNA.3.1 empty vector. Cells were harvested 24 h post-transfection and WCLs analysed by western blotting.

## Acknowledgements

We thank Professor Juergen Hass (University of Edinburgh) and Dr Benjamin Hale (University of Zurich), Professor Roger Everett (MRC-UoG CVR) for the provision of reagents, and Dr Seema Jasim (University of Edinburgh) for experimental assistance.

## Supplemental Figure Legends

**S1 FIG. TRIM22 is constitutively expressed to high level in lung tissue.** (A) Transcript expression levels (log2 median TPM) of 67 TRIM family members in human lung tissue biopsies (n=457; S1A Table). Black line: median TRIM transcript expression; whisker: 5^th^ to 95^th^ percentile range. (B) Transcript expression levels (log2 median TPM) of TRIM22 across a range of human tissues (S1C Table). Lung and spleen tissues are highlighted (red and blue circles, respectively). Black line: median; whisker: 5^th^ to 95^th^ percentile range. (C, D) Violin plots showing individual TRIM family member (C, lung) or TRIM22 (D, all tissues) expression profiles (log10 TPM), respectively. White line; median. Box; 5^th^ and 95^th^ percentile range. (A-D) Data adapted under creative commons license from GTEx portal [64].

**S2 FIG. TRIM22 antibody validation.** MRC5t cells were stably transduced to express non-targeting control (shCtrl) or TRIM22-targeting (shTRIM22, clones B7-B9) shRNAs. (A) qRT-PCR quantitation of *TRIM22* mRNA levels in MRC5t shCtrl and shTRIM22 cells. Values normalized to shCtrl. Mean RQ and RQ min/max shown. (B) Western blot of WCLs derived from MRC5t shCtrl and shTRIM22 (clone B7) cells showing TRIM22 (detected using pAb HPA003575; Sigma-Aldrich) expression levels. Actin is shown as a loading control. (C) Confocal micrographs showing the nuclear localization of TRIM22 in MRC5t shCtrl or shTRIM22 (clone B7) cells. TRIM22 was detected by indirect immunofluorescence (pAb HPA003575; Sigma-Aldrich). Nuclei were stained with DAPI.

**S3 FIG. Constitutive TRIM22 expression is lost in many transformed cell lines.** MRC5 (primary lung fibroblast), MRC5t (telomerase immortalized MRC5), A549, HEK 293T (293T), HeLa, and HEp2 cells were treated for 24 h with (+) or without (-) IFN-β stimulation (100 IU/ml). (A) qRT-PCR quantitation of *TRIM22* mRNA transcript levels across the panel of cell lines. Values normalized to *TRIM22* levels in MRC5t cells without IFN-β stimulation. n=3, means and SD shown. (B) Western blots showing TRIM22 expression levels in WCLs derived from the panel cell lines. Actin is shown as a loading control.

**S4 FIG. Identification of tissue-specific and cell-type dependent patterns of constitutive ISG expression.** (A) Transcript expression profiles (log2 TPM) of 178 ISGs in HBEC3, A549, HEK 293 (293), and HeLa cells plotted against the equivalent gene set from human lung tissue biopsies (n=427; log2 median TPM shown). Every second gene labelled (S3A Table). (B) Violin plot showing cell line and lung tissue ISG expression profiles (as in A). Horizontal solid lines; median ISG expression. Horizontal dotted lines; 5^th^ and 95^th^ percentile range. Paired one-way ANOVA (Friedman multiple comparison test); **** *P* < 0.0001. (C) Heatmap showing clustered distribution of ISG transcript levels between cell lines and human lung tissue (as in B). Cell line and tissue RNA-seq data adapted under creative commons licence from HPA and GTEx portal, respectively [62–64].

**S5 FIG. Annotated STRING network of downregulated immune system genes in virally-transformed cells.** Annotated high-confidence (> 0.7) STRING network of identified differential downregulated immune system genes in virally-transformed cells (as shown in Fig 9C; S4D Table). Downregulated ISGs common to three transformed cell lines; core, blue circles: common to two transformed cell lines; shared, purple circles: unique to one transformed cell line; unique, grey circles. Network PPI enrichment: *P* < 1.0e-16. Ranked biological process (GO) and KEGG pathways with associated counts in gene sets (count) and FDR (false discovery rate) values shown.

**S6 FIG. TRIM family expression profile in human lung tissue and lung epithelial (HBEC3 and A549) cells.** (A) Transcript expression profile of 67 TRIM family members derived from human lung tissue, hTERT immortalized (HBEC3), or virally-transformed (A549) human lung epithelial cells. Data adapted under creative commons license from Human Protein Atlas (HPA; [62, 63]) and Genotype-Tissue Expression (GTEx; [64]) project. (B) Scatter plots showing the differential transcript expression of individual TRIM family members between lung tissue data sets (GTEx and HPA), lung tissue (GTEx) and either HBEC3 or A549, or between HBEC3 and A549 (as indicated). Solid coloured lines; linear regression. Solid black lines; 95% confidence interval. R-squared (R^2^) values indicated. Selected TRIMs are highlighted for reference.

## Supplemental Tables

**S1 Tables. TRIM family member transcript expression values across a range of human tissues.** (S1A) HPA and GTEx transcript expression values of 67 TRIM family member genes in human lung tissue. (S1B) HPA transcript expression values of TRIM22, TRIM32, and TRIM41 across a range of human tissues (as indicated). (S1C) GTEx TRIM22 transcript expression values across a range of human tissues (as indicated). Data adapted under creative commons license from Human Protein Atlas (HPA; [62, 63]) and Genotype-Tissue Expression (GTEx; [64]) project.

**S2 Tables. ISG transcript expression values across a range of human tissues.** (S2A) Transcript expression values for 200 ISGs (previously shown to be upregulated ≥ 8-fold change in response to universal IFN treatment in primary cell culture; [67]) across a range of mucosal (lung, small intestine [int.; terminal ileum], esophagus [mucosa], colon [sigmoid]), and non-mucosal (liver, skin [suprapubic], and kidney [cortex] tissues. (S2B) Transcript expression values of IFN-related receptors and cytokines across human tissues (as in S2A). Data adapted under creative commons license from Genotype-Tissue Expression (GTEx; [64]) project.

**S3 Tables. Constitutive ISG transcript expression values across a range of cell lines and lung tissue.** (S3A) Transcript expression values for 178 ISGs (previously shown to be upregulated ≥ 8-fold change in response to universal IFN treatment in primary cell culture; [67]) across a range of non-treated transformed (A549, HEK 293, and HeLa) cells, hTERT immortalized (HBEC3) cells, or lung tissue. (S3B) ISG expression values of differentially downregulated ISGs between cell lines (HBEC3 ≥ 5-fold change, ≥ 5 TPM) and equivalent gene set tissue expression values obtained from human lung tissue biopsies (n=427, log2 median TPM). Data adapted under creative commons license from Human Protein Atlas (HPA; [62, 63]) and Genotype-Tissue Expression (GTEx; [64]) project.

**S4 Tables. Constitutive immune system transcript expression values across a range of cell lines.** (S4A) Transcript expression values for differentially downregulated genes identified between non-treated HBEC3 (≥ 3 TPM [median TPM all genes]) and A549 [median 3 TPM all genes], HEK 293 [median 3 TPM all genes], or HeLa [median 4 TPM all genes] cells (≤ 4 TPM; HBEC3 ≥ 5-fold change; blue circles in Fig 9A). (S4B) Reactome (https://reactome.org) pathway analysis of downregulated mapped genes (as in S4A; blue circles in Fig 9A). Red text highlights pathways relating to the immune system. (S4C) Uniprot IDs for downregulated genes mapped to the immune system (as in B; Reactome immune system). (S4D) Ranked transcript expression values for downregulated immune system genes identified in all three (core; blue), common to two (shared; purple), or unique to one (unique) transformed cell line(s). Data adapted under creative commons license from Human Protein Atlas (HPA; [62, 63]).

**S5 Tables. Extended IAV KEGG network transcript expression values across a range of cell lines and lung tissue.** (S5A) Transcript expression values of 184 IAV immuno-regulatory genes (extended (ext.) IAV KEGG network (<u>hsa05164</u>), [70]) in non-treated HBEC3, A549, HEK 293, and HeLa cells. (S5B) Transcript expression values of differentially downregulated IAV immuno-regulatory genes (HBEC3 ≥ 5-fold change, ≥ 3 TPM) and equivalent gene set expression values obtained from human lung tissue biopsies (n=427; log2 median TPM). Data adapted under creative commons license from Human Protein Atlas (HPA; [62, 63]) and Genotype-Tissue Expression (GTEx; [64]) project.

**S6 Table. TRIM family member transcript expression values across a range of cell lines and lung tissue.** Transcript expression values of 67 TRIM family member genes across a range of non-treated cells (HBEC3, A549, HEK 293, and HeLa) and human lung tissue biopsies (n=427; log2 median TPM). Data adapted under creative commons license from Human Protein Atlas (HPA; [62, 63]) and Genotype-Tissue Expression (GTEx; [64]) project.

**S7 Table. Custom IAV primer-probe sequences.** Nucleotide sequences for custom IAV (*NP*, *M1*, *NS1*/*NEP*) primer-probes mixes used in the study.

